# Neuroinflammation plays a critical role in cerebral cavernous malformation disease

**DOI:** 10.1101/2022.05.09.491214

**Authors:** Catherine Chinhchu Lai, Bliss Nelsen, Eduardo Frias-Anaya, Helios Gallego-Gutierrez, Marco Orecchioni, Hao Sun, Omar A. Mesarwi, Klaus Ley, Brendan Gongol, Miguel Alejandro Lopez-Ramirez

## Abstract

**Background:** Cerebral Cavernous Malformations (CCMs) are neurovascular lesions caused by loss-of-function mutations in one of three genes, including KRIT1 (CCM1), CCM2, and PDCD10 (CCM3). CCMs affect ∼1/200 children and adults, and no pharmacologic therapy is available. CCM lesion count, size, and aggressiveness vary widely among patients of similar ages with the same mutation or even within members of the same family. However, what determines the transition from quiescent lesions into mature and active (aggressive) CCM lesions is unknown.

**Methods:** We use genetic, RNA-seq, histology, flow cytometry and imaging techniques to report the interaction between CCM-endothelium, astrocytes, leukocytes, microglia/macrophages, neutrophils (CALMN interaction) during the pathogenesis of CCMs in the brain tissue.

**Results:** Expression profile of astrocytes in adult mouse brains using translated mRNAs obtained from the purification of EGFP-tagged ribosomes (*Aldh1l1-EGFP/Rpl10a*) in the presence or absence of CCM lesions (*Slco1c1-iCreERT2;Pdcd10^fl/fl^*; *Pdcd10^BECKO^*) identifies a novel gene signature for neuroinflammatory astrogliosis. CCM reactive astrocytes have a neuroinflammatory capacity by expressing genes involved in angiogenesis, chemotaxis, hypoxia signaling, and inflammation. RNA-seq analysis on RNA isolated from brain endothelial cells (BECs) in chronic *Pdcd10^BECKO^* mice (CCM-endothelium), identified crucial genes involved in recruiting inflammatory cells and thrombus formation through chemotaxis and coagulation pathways. In addition, CCM- endothelium was associated with increased expression of *Nlrp3* and *Il1b*. Pharmacological inhibition of NLRP3 significantly decreased inflammasome activity as assessed by quantification of a fluorescent indicator of caspase-1 activity (FAM-FLICA caspase-1) in BECs from *Pdcd10^BECKO^* in chronic stage. Importantly, our results support the hypothesis of the crosstalk between astrocytes and CCM endothelium that can trigger recruitment of inflammatory cells arising from brain parenchyma (microglia) and the peripheral immune system (leukocytes) into mature active CCM lesions that propagate lesion growth, immunothrombosis, and bleedings. Unexpectedly, partial or total loss of brain endothelial NF-kB activity (using *Ikkb^fl/fl^* mice) in chronic *Pdcd10^BECKO^* mice does not prevent lesion genesis or neuroinflammation. Instead, this resulted in elevated number of lesions and immunothrombosis, suggesting that therapeutic approaches designed to target inflammation through endothelial NF-kB inhibition may contribute to detrimental side effects.

**Conclusions:** Our study reveals previously unknown links between neuroinflammatory astrocytes and inflamed CCM endothelium as contributors that trigger leukocyte recruitment and precipitate immunothrombosis in CCM lesions. However, therapeutic approaches targeting brain endothelial NF-kB activity may contribute to detrimental side effects.

## Introduction

Cerebral cavernous malformations (CCMs) are common neurovascular lesions causing a lifelong risk of brain hemorrhage, seizures, and neurological sequelae for which there is no current effective pharmacologic therapy ^1, 2^. CCMs affect approximately 0.5% of the general population, in which children represent ∼25% of diagnosed individuals ^3, 4^. Inherited germline (∼20%) and somatic (∼80%) loss of function mutations in the genes *KRIT1* (Krev1 interaction trapped gene 1, CCM1), *CCM2* (Malcavernin), *PDCD10* (programmed cell death protein 10, CCM3) propel brain vascular changes marked by the disruption of intercellular junctions, increase in reactive oxygen species (ROS), angiogenesis, altered basement membrane composition, and increased vascular permeability ^2, 5–10^. CCMs are dynamic lesions that can form, enlarge, regress, or behave aggressively, producing repetitive hemorrhage that contributes to the clinical manifestation of CCM disease in familial and sporadic cases ^11–13^. Moreover, histological analysis in human CCM lesions has suggested that CCM bleeding in brain tissue may lead to inflammation associated with adaptive immune response and thrombosis in varying degrees ^14–16^.

Importantly, histopathological hallmarks identified in human CCM brain tissue are closely recapitulated in the central nervous system (CNS) of genetically sensitized CCM mouse models ^17, 18^ and, more recently observed, in chronic CCM mouse models using inducible brain endothelial-specific genetic inactivation of CCM genes ^19–23^. These CCM animal models have been crucial to unveiling transition phases in CCM lesion genesis that range from early-stage isolated caverns (stage 1) during CCM lesion formation to late-stage multi-cavernous lesions (stage 2) containing hemosiderin deposits and immune cell infiltration (characteristics of human active CCM lesions) (18). However, the molecular and cellular mechanisms contributing to CCM lesion initiation and progression as well as the transition between quiescence to mature active lesions in the CNS remains elusive (24). Recent studies indicate that GFAP+ astrocytes contribute to CCM pathogenesis by integrating a circuit of neurovascular dysfunction during CCM lesion formation ^22^. These findings suggest hypoxia programs are activated during CCM formation that affects endothelial-astrocyte crosstalk associated with changes in angiogenesis, inflammation, and endothelial-cell metabolism, involved in CCM formation.

The present study shows that while the hypoxia and angiogenesis pathways are important in CCM lesion formation^22^, the inflammation and NLRP3 inflammasome activity play a more critical role in transitioning into mature and active CCM lesions. We report that neuroinflammation conferred by neuroinflammatory astrogliosis and CCM endothelium attracts innate and adaptive immunity to CCM lesions in a chronic and inducible CCM mouse models. Unexpectedly, partial or total loss of brain endothelial NF-kB activity in a chronic CCM animal model does not prevent lesion genesis or neuroinflammation. Instead, this loss results in elevated immunothrombosis, suggesting that therapeutic approaches designed to target inflammation through NF-kB inhibition may contribute to detrimental side effects. Here, we propose that the interaction between CCM-endothelium, astrocytes, leukocytes, microglia/macrophages, neutrophils (CALMN interaction) play a critical role in the pathogenesis of CCMs by inducing active lesions and propensity to immunothrombosis. Our study reveals a new role for astrocytes and CCM endothelium in neuroinflammation and identifies that “CALMN interaction”, may lead to new therapeutic strategies to prevent active CCM lesions.

## Results

### Increase of hypoxia, inflammation and inflammasome signaling pathways in CCM disease

Histological analysis of CCM lesions in chronic animal models has found that lesions vary from early-stage isolated caverns (stage 1) that range of size to late-stage multicavernous lesions (stage 2) more resembling of human CCM lesions^17^. However, the molecular and cellular mechanism contributing to CCM lesion initiation and activation is unknown ^24^. Therefore, to monitor changes in CCM lesion genesis, conferred exclusively to the CNS tissue, at different developmental stages, we created a brain microvascular endothelial cell-specific *Pdcd10* knock out animal model (*Pdcd10^BECKO^*) by crossing a brain endothelial tamoxifen-regulated Cre (*Slco1c1-CreERT2*) with *Pdcd10^fl/fl^* mice (*Slco1c1-CreERT2; Pdcd10^fl/fl^*) ^19, 20, 22^ (Fig. 1A). We have chosen to study the CCM lesions in the cerebrum because it offers a temporal pattern of lesion genesis representative to the CNS. Upon tamoxifen injection, the brains of *Pdcd10^BECKO^* animals develop CCM lesions progressively that are subdivided into acute (P15), progression (P50), and chronic (P80) stages (Fig. 1A)^17, 19, 22^. Moreover, differences in cerebrum histological changes correlated with changes in gene expression pattern in cerebral tissue (Fig 1A, 1B) that coincided with the induction of acute phase CCM marker genes including the HIF-target genes *Loxl2,* and *Vegfa* ^22^(Fig. 1B, Supplemental Fig 1). We observed a marked elevation in the expression of *Loxl2* and *Angpl4* in the progression and chronic stages. However, the levels of *Vegfa* mRNA were attenuated in chronic phase CCM brain tissue (Fig. 1B), suggesting that angiogenesis triggered by VEGF is critical in CCM lesion formation. Furthermore, we observed a significant increase of neuroinflammatory genes in the CCM cerebral tissue, including*, Mcp1* and *Cd74*, as well as inflammasome genes, such as *Il1b* and *Nlrp3,* duringin the progression and chronic stages (Fig. 1B, Supplemental Fig 1). We also observed that among genes associated with coagulation pathways in CCM, *Procr* (EPCR) ^23, 25, 26^ is the best candidate gene to differentiate between disease severities (Supplemental Fig 1). These results suggest that while hypoxia and angiogenesis signaling are important in the CCM lesion formation^22^, neuroinflammation signaling may play a more important role in the progression and chronic stages. In agreement with previous results, prominent GFAP-immunoreactivity was observed in astrocytes flanking acute and chronic cerebral CCM lesions^22^ (Fig. 1C, 1D). Collectively, these results suggest that hypoxia, neuroinflammation, and inflammasome signaling are present in mature CCM lesions and that astrogliosis is a critical feature in the disease.

**Fig.1.**
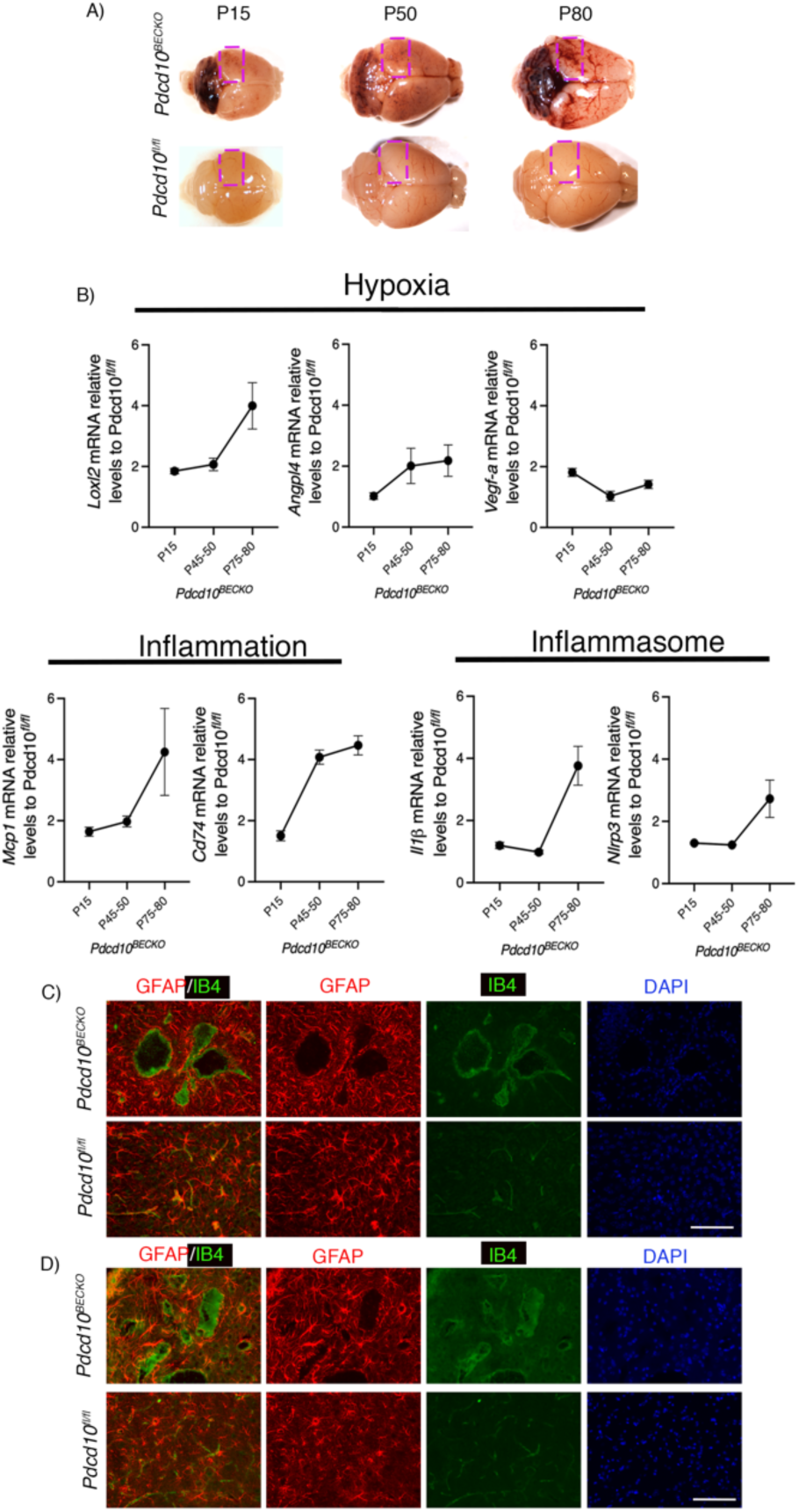
Increase of hypoxia and neuroinflammation signaling pathways in CCM disease. **A,** CCM lesions are present in the cerebellum and cerebrum of *Pdcd10^BECKO^* mice at acute (P15), progression (P50), and chronic stage (P80). A square shows the region used in *B* for RNA analysis. **B,** Analysis of *Loxl2*, *Angpl4*, *Vegfa*, *Mcp1*, *Cd74*, *Il1β*, and *Nlrp3* mRNA levels by RT-qPCR in cerebral tissue from mice at acute, progression, and chronic stage as indicated. Data are mean ±SEM, n = 3 or 6 mice in each group. **C,** Immunofluorescence staining of GFAP+ astrocytes (red), and endothelial marker isolectin B4 (IB4; green) of cerebral sections from P15 *Pdcd10^BECKO^* and littermate control *Pdcd10^flfl^*. DAPI staining (blue) was used to reveal nuclei. n = 4 mice in each group. Scale bar, 100 µm. **D,** Immunofluorescence staining of GFAP+ astrocytes (red), and endothelial marker isolectin B4 (IB4; green) of cerebral sections from P80 *Pdcd10^BECKO^* and littermate control *Pdcd10^flfl^*. DAPI staining (blue) was used to reveal nuclei. n = 5 mice in each group. Scale bar, 100 µm.

### CCM endothelium induces neuroinflammatory astrocytes

To better understand how GFAP+ astrocytes contribute to CCM pathogenesis, the astrocyte ribosome-bound mRNA levels were analyzed using Translational Ribosome Affinity Purification (TRAP) as a measurement of active translation ^27, 28^. For these experiments, *Pdcd10^BECKO^* and *Pdcd10^fl/fl^* control mice were crossed with a ribosome tagged mouse line (*Aldh1l1-EGFP/Rpl10a)*. Following CCM induction with tamoxifen administration, the level of ribosome-bound and global mRNA abundance was quantified (Fig. 2). The presence of GFAP+ astrocytes was observed along with EGFP-RpL10a co-localization in *Pdcd10^fl/fl^;Aldh1l1-EGFP/Rpl10a* brain sections. We also observed that CCM lesions in *Pdcd10^BECKO^;Aldh1l1-EGFP/Rpl10a* develop surrounded by GFAP+ astrocytes that colocalized with EGFP-RpL10a expressing cells (Fig. 2A, Supplemental Fig. 2). Ribosome-bound mRNAs were highly enriched for astrocyte-specific genes and almost absent of other brain resident cell markers (Supplemental Fig. 2). We identified 153 upregulated and 168 downregulated genes (FDR < 0.03) between astrocyte TRAP in *Pdcd10^BECKO^;Aldh1l1-EGFP/Rpl10a* and astrocyte TRAP in *Pdcd10^fl/fl^;Aldh1l1-EGFP/Rpl10a* (Fig. 2B, Supplemental Fig. 3). Gene functional classification analysis ^29^ of differentially expressed genes (DEGs) revealed significant enrichment for terms related to antigen presentation, platelet activation/aggregation, VEGF signaling, and glucose metabolism (Fig. 2C). Genes demonstrating the most significant fold change include genes associated with neurodegenerative diseases (e.g., *Ndufa4l2*, *App*, *S1pr3*) (Fig. 2D). Moreover, many relevant DEGs were also associated with hypoxia *(e.g., Adm*, *Igfbp3*, *Loxl2*), chemokines (e.g., *Cx3cl1, Ccl2, Ackr1, Cx3cr1*), and inflammation (e.g., *Cd74, Adora2a, C1qa*) (Fig. 2e,2f,2g). Of particular interest are *Cx3cl1* and *Ccl2* which enable the capacity of reactive astrocytes to recruit microglia and leukocytes to CCM lesions ^30, 31^. Indeed, inflammation, neurodegeneration, and ischemia are CNS injuries that induce astrocyte polarization into a neuroinflammatory A1 or neuroprotective A2 reactive state^32–34^. However, we observed no clear polarization of astrocytes during CCM disease toward A1-like or A2-like reactive astrocytes (Fig. 2H, Supplemental Fig. 3). Instead, we observed that astrocytes activated during CCM disease acquire a particular transcriptional signature that we denominated A3 CCM reactive astrocyte with neuroinflammatory capacity (Fig. 2H, 2G, Supplemental Fig. 3). These results indicate a crucial role for a subset of neuroinflammatory reactive astrocytes in recruiting inflammatory cells to CCM mature lesions.

**Fig. 2.**
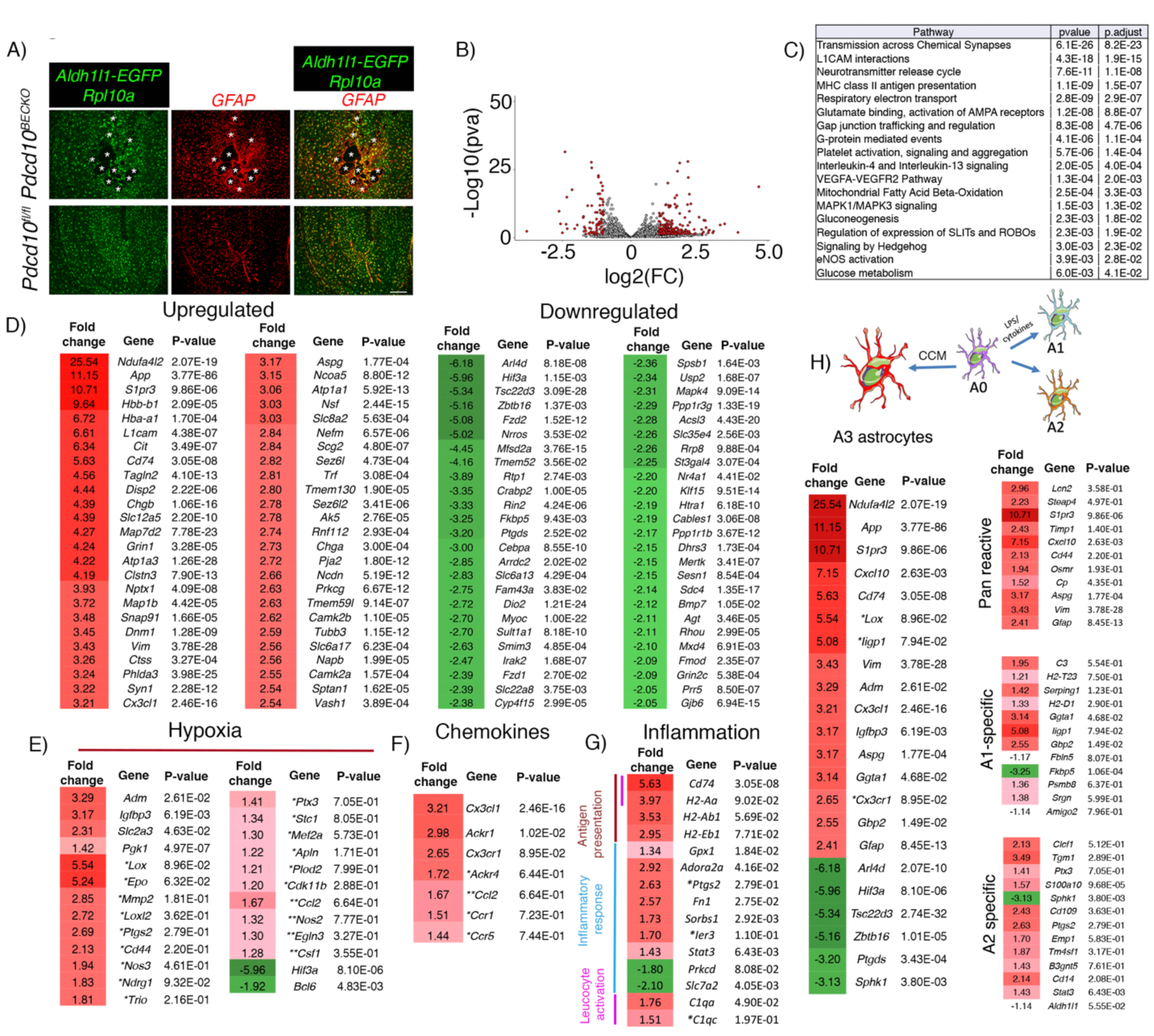
Neuroinflammatory astrocytes in CCM disease. **A,** Immunofluorescence analysis show colocalization of GFAP+ astrocytes with cells expressing EGFP-RpL10a in *Pdcd10^fl/fl^;Aldh1l1-EGFP/Rpl10a* brains sections. CCM lesions in *Pdcd10^BECKO^;Aldh1l1-EGFP/Rpl10a* develop surrounded by GFAP+ astrocytes that colocalized with EGFP-RpL10a expressing cells. Asterisks denote the vascular lumen of CCM lesions. Scale bar is 200 µm. n = 3 mice in each group. **B,** Volcano plot of differentially expressed transcripts in P75 *Pdcd10^BECKO^;Aldh1l1-EGFP/Rpl10a* versus littermate control *Pdcd10^fl/fl^;Aldh1l1-EGFP/Rpl10a.* Transcripts are represented on a log2 scale. The significantly down- and up-regulated genes are labeled in red. n = 3 mice in each group. **C,** Top pathway enrichment analysis for *Pdcd10^BECKO^;Aldh1l1-EGFP/Rpl10a* brain tissue. *Pdcd10^fl/fl^;Aldh1l1-EGFP/Rpl10a* littermates were used as controls. n = 3 mice in each group. (320 DEG, fold change ≥ 1.2 and FDR < 0.03). **D,** List of the top 50 up- or down-regulated genes from translated mRNAs obtained from ribosomes in astrocytes from *Pdcd10^BECKO^;Aldh1l1-EGFP/Rpl10a* brains. Fold change and *P* values are shown for each gene. n = 3 mice in each group. **E,** Analysis of hypoxia-regulated genes from *Pdcd10^BECKO^;Aldh1l1-EGFP/Rpl10a* brains following astrocyte-TRAP-isolated RNA. **F,** Analysis of chemokine transcripts from *Pdcd10^BECKO^;Aldh1l1-EGFP/Rpl10a* brains following astrocyte-TRAP-isolated RNA. **G,** Analysis of inflammation-related transcripts from *Pdcd10^BECKO^;Aldh1l1-EGFP/Rpl10a* brains following astrocyte-TRAP-isolated RNA. **H,** Transcriptional signature of neuroinflammatory 3A astrocytes in CCM disease. Pan reactive, A1 specific and A2 specific genes are shown. Fold change and *P* values are shown for each gene. n = 3 mice in each group. * and ** indicate genes that do not reach statistical significance because one of the biological replicates the fold change was too large but in the same direction (*) or when one of the biological replicates does not change (**).

### Increase of hypoxia, inflammation and inflammasome signaling pathways in CCM endothelium

CCM endothelium induces HIF-1α protein stabilization in astrocytes under normoxic conditions through elevation of nitric oxide, resulting in the synthesis of astrocyte-derived VEGF and enhancement of CCM lesion formation ^22^. To better understand the interaction between CCM endothelium and astrocytes and potentially other neurovascular cells during CCM disease, we performed an RNA-seq profiling of brain endothelial cells (BECs) isolated from P75 *Pdcd10^BECKO^;Aldh1l1-EGFP/Rpl10a* brains and *Pdcd10^fl/fl^;Aldh1l1-EGFP/Rpl10a* control brains (Fig. 3A). We identified 827 upregulated and 526 downregulated genes between BECs from *Pdcd10^BECKO^* and *Pdcd10^fl/fl^* controls (Fig. 3A, Supplemental Fig.4). The mRNAs obtained from the purification of BECs were highly enriched for brain endothelial-specific genes, but a small increase in leukocyte and myelin gene contaminants was also detected (Fig. 3B). Gene set enrichment analysis (GSEA) of DEGs in BECs isolated from *Pdcd10^BECKO^* brains revealed significant enrichment for terms related to neutrophil degranulation, platelet activation/aggregation, VEGF, ROS, ROCK, and EndMT signaling (Fig. 3C). The transcriptomic analysis also revealed significant upregulation of additional pathways associated with aneurysms, neurodegenerative diseases, and senescence genes (e.g., *Col10a1*, *Serpina1e/b/d*, *Cyp3a13, Gkn3, Serpine1*). Moreover, hypoxia genes (e.g., *Apln*, *Egln3*, *Vegfa*, *Loxl2*, *Cd44*) were also predominantly upregulated in BECs during chronic CCM disease (Fig. 3E). Importantly, genes implicated in neuroinflammatory response (e.g., *Il6*, *Ccl2*, *Cd74, Ccl2, Cx3cl1, Ackr2*) and inflammasome activity (e.g., *Il1b*, *Nlrp3*) underscored a potential role of CCM endothelium-induced neuroinflammation (Fig. 3F through 3H). Of note, CCM endothelium is associated with locally increased expression of anticoagulant endothelial receptors that create a local bleeding diathesis in CCMs^25^. We observed a significant upregulation of genes implicated in the balance between the anti- and pro-coagulation system (e.g., *Procr* (EPCR), *Thbd*, *Serpine1* (PAI-1), *Tfpi* (TFPI), *Vwf*) (Fig. 3H). These results suggest that anti- and pro-thrombotic factors co-express on mature CCM lesions. Moreover, as previously reported, we observe a significant increase in genes associated with ROS in the CCM endothelium (e.g., *Cybb*, *Slc7a2*, *Ncf4*) (Fig. 3h)^35^. These transcriptomic analyses from CCM endothelium and astrocytes are consistent with our immunohistology, and bulk RNA gene expression analyses initially observed in the chronic CCM brains (Fig. 1) and suggest a complex cellular interaction occurring during CCM lesion maturation where hypoxia and neuroinflammation signaling pathways from both CCM endothelium and astrocytes are critical.

**Fig. 3.**
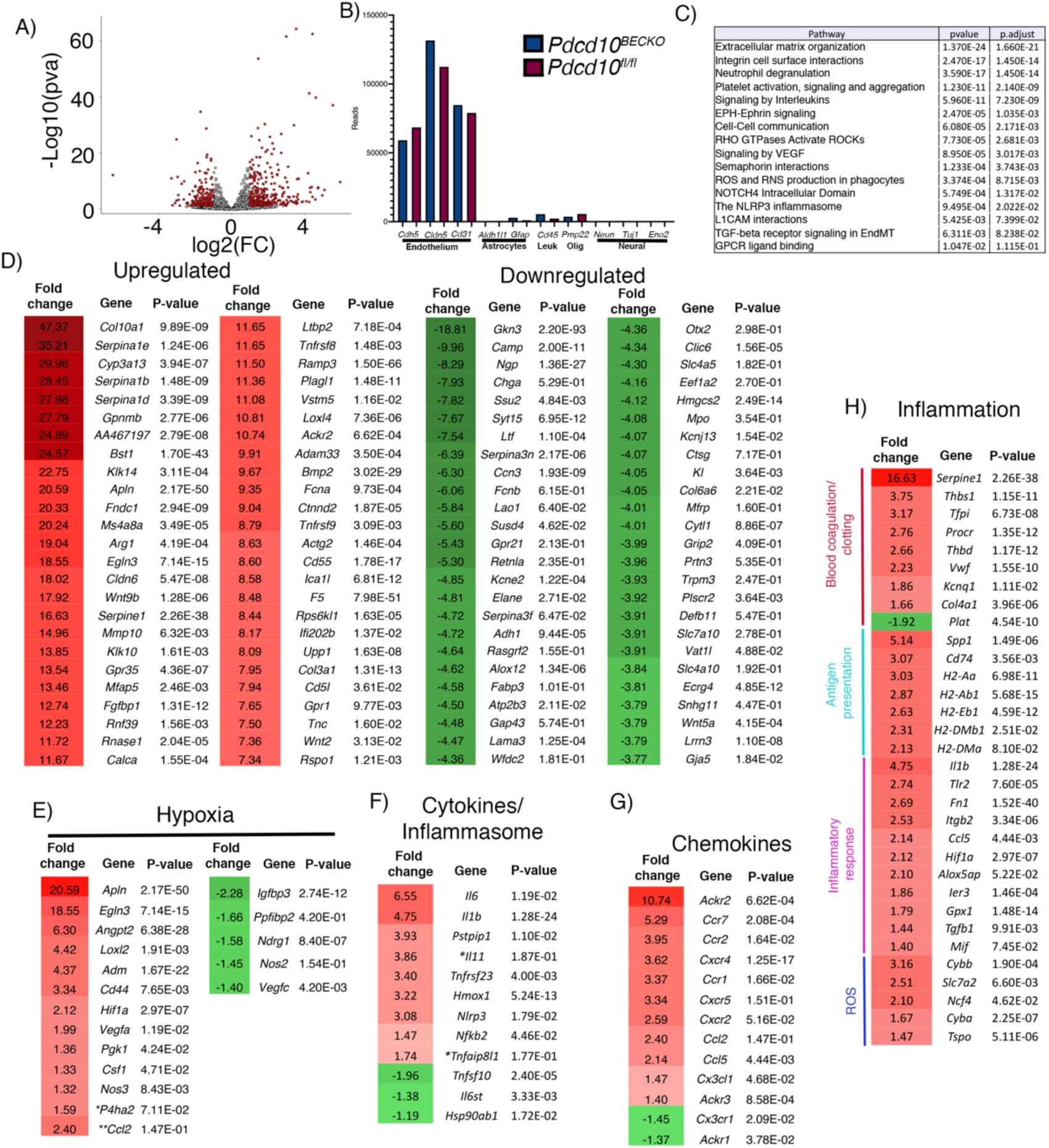
Inflammation and inflammasome pathways are increased in the CCM endothelium. **A,** Volcano plot of differentially expressed transcripts in fresh isolated brain endothelial cells from P75 *Pdcd10^BECKO^;Aldh1l1-EGFP/Rpl10a* versus isolated brain endothelial cells from littermate control *Pdcd10^fl/fl^;Aldh1l1-EGFP/Rpl10a.* Transcripts are represented on a log2 scale. The significantly down- and up-regulated genes are labeled in red. n = 3 or 4 mice in each group. **B,** Analysis of known brain cell-specific gene markers from *Pdcd10^BECKO^;Aldh1l1-EGFP/Rpl10a* (blue bar) and *Pdcd10^fl/fl^;Aldh1l1-EGFP/Rpl10a* littermate controls (red bar) following isolated brain endothelial cells. n = 3 or 4 mice in each group. **C,** Top pathway enrichment analysis for *Pdcd10^BECKO^;Aldh1l1-EGFP/Rpl10a* isolated brain endothelial cells. n = 3 or 4 mice in each group. **D,** List of the top 50 up- or down-regulated genes from isolated brain endothelial cells obtained from *Pdcd10^BECKO^;Aldh1l1-EGFP/Rpl10a* brains. Fold change and *P* values are shown for each gene. n = 3 or 4 mice in each group. **E,** Analysis of hypoxia-regulated genes from *Pdcd10^BECKO^;Aldh1l1-EGFP/Rpl10a* isolated brain endothelial cells. **F,** Analysis of cytokines and inflammasome transcripts from *Pdcd10^BECKO^;Aldh1l1-EGFP/Rpl10a* isolated brain endothelial cells. **G,** Analysis of chemokine transcripts from *Pdcd10^BECKO^;Aldh1l1-EGFP/Rpl10a* isolated brain endothelial cells. **H,** Analysis of inflammation-related transcripts from *Pdcd10^BECKO^;Aldh1l1-EGFP/Rpl10a* isolated brain endothelial cells. Fold change and *P* values are shown for each gene. n = 3 or 4 mice in each group.

### Augmented inflammasome activity in CCM endothelium

During inflammation, the transcription factor nuclear factor *k*B (NF−*k*b) is critical to upregulate NLRP3 gene expression ^36^. NLRP3 inflammasome assembly and activation in endothelial cells is critical in developing and progressing chronic and neurodegenerative diseases ^36–38^. Therefore, because inflammasome activation in the CCM endothelium may be an important component of the neuroinflammation observed in CCM disease, we aimed to investigate inflammasome activity in the chronic CCM mouse animal model. Caspase-1 activity is controlled by the NLRP3 inflammasome and a fluorescent indicator of caspase-1 activity (FAM-FLICA caspase-1) allows us to quantitatively assess the inflammasome activity in isolated BECs from P75 *Pdcd10^BECKO^* brains (Fig. 4A through 4C). We observed that caspase-1 activation was significantly increased in subsets of isolated BECs from *Pdcd10^BECKO^* brains when compared to BECs from *Pdcd10^fl/fl^* brains (Fig. 4A, 4B). We confirmed the specificity of BECs to elevate FAM-FLICA staining in *Pdcd10^BECKO^* brains by in situ treatment with MCC950, an inhibitor of the NLRP3 inflammasome activation ^39^ that significantly decreases the FAM-FLICA staining in BECs from *Pdcd10^BECKO^* when compared with same BECs treated with vehicle (Fig. 4A, 4C). Moreover, the use of CCM brain tissue sections confirmed caspase1 activation in the brain endothelium forming multicavernous lesions (stage 2), and in leukocytes located within the multicavernous (Fig. 4D). However, we did not observe an obvious caspase1 activation in single CCM lesions (Data not shown). These results suggest that the chronicity and the presence of inflammatory leukocytes contribute to inflammasome activation in CCM lesions. Since NF−*k*b activity is essential for NLRP3 expression, inflammasome priming, and assembly ^40, 41^, we next investigated whether RelA (p65) levels were altered in CCM brains. Levels of p65 protein were increased in *Pdcd10^BECKO^* brains in comparison with littermate control *Pdcd10^fl/fl^* brains (Fig. 4E). This data revealed that mature CCM lesions are poised to innate immune response and subsequent immunothrombosis by elevation of brain endothelial inflammasome activity.

**Fig. 4.**
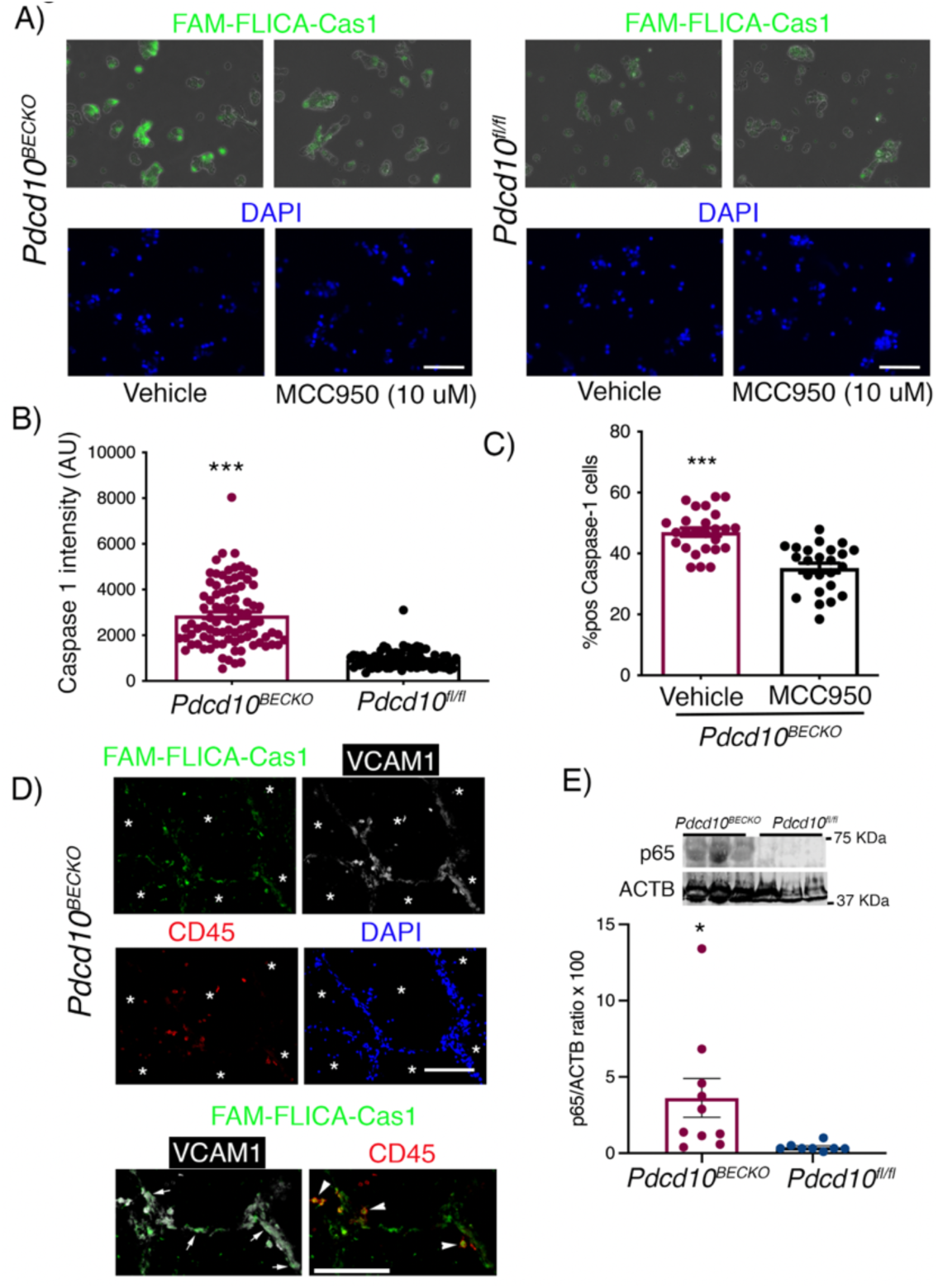
Increase in inflammasome activity in CCM endothelium. **A,** Visualization of inflammasome activation in P80 *Pdcd10^BECKO^* isolated brain endothelial cells in the presence or absence of MCC950 (10 µM, 1 h at 37 C). *Pdcd10^fl/fl^* isolated brain endothelial cells were used as controls in the presence or absence of MCC950. FAM-FLICA Caspase-1 assay kit was used to visualize caspase-1 signal as an indicator of inflammasome activity. Scale bars, 50 µm. **B,** Quantification of FAM-FLICA Caspase-1 intensity from *a* in *Pdcd10^BECKO^* compared to *Pdcd10^fl/fl^* animals. **C,** Percentage of caspase-1 positive *Pdcd10^BECKO^* isolated cells exposed to MCC950 compared to vehicle. Data are mean ±SEM n = 2 or 3 mice in each group. **D,** FAM-FLICA fluorescence analysis (green) in multi cavernous CCM lesions in combination with immunofluorescence for VCAM-1 (white) and CD45 (red), and DAPI labelling nuclei (blue). Asterisks denote vascular lumen of lesions. FAM-FLICA Caspase-1 signal is observed in both VCAM-1 (endothelium) and CD45 (leukocytes) positive cells as marked by white arrows and arrowheads, respectively. Scale bars, 100 µm. **E,** Quantification of p65 protein in P80 *Pdcd10^BECKO^* brains compared to *Pdcd10^fl/fl^* brain controls by western blot. Data are mean ±SEM n = 8 or 10 mice in each group. *, *P*<0.05, ***, *P*<0.001.

### Increase of immune cells in CCM brain tissue

We next evaluated whether CCM endothelial-astrocyte neuroinflammatory signaling during CCM disease prompts the recruitment of immune cells into the CCM brain tissue. As anticipated, we observed significant recruitment of CD45+ cells as well as early vascular thrombosis (staining for CD41+ platelet aggregation) in the vascular lumen of lesions in P75 *Pdcd10^BECKO^* brains (Fig. 5A). In agreement with reports from human CCM lesions ^16, 42^, histological analysis of P75 *Pdcd10^BECKO^* brains also showed large mature vascular thrombosis associated with CD45+ leukocyte infiltration (Fig. 5B, 5C), and in a serial section, show the accumulation of neutrophils lining the vessel wall and within the thrombus (Supplemental Fig. 5, Supplemental Fig. 6). Histological analysis of CCM brain lesions also demonstrates that CX3CR1+ and CD16/32+ cells constitute the larger number of cells forming the vascular thrombosis (Supplemental Fig. 6). We used flow cytometry to further determine the identity and quantification of brain-infiltrating leukocytes during CCM disease. Consistent with histological analysis, we first observed a significant increase in CD45+ leukocytes isolated from P75 *Pdcd10^BECKO^* brains compared to leukocytes isolated from *Pdcd10^fl/fl^* control brains (Fig. 5D). Flow cytometry results show a significant increase in infiltrating myeloid cells in *Pdcd10^BECKO^* brains (Fig. 5E). We identified classical monocytes as CX3CR1-CD11b+Ly6C+, and microglia as CX3CR1+CD11b+F4/80+. The phenotype Ly6G+CD11b+, CX3CR1+CD11b+F4/80- Ly6C-, and CD11b+CX3CR1-Ly6C-CD11c+ identified neutrophils, non-classical monocytes, and dendritic cells, respectively (Fig 5E). Moreover, we further observed that lymphoid cells were elevated in *Pdcd10^BECKO^* brains (Fig. 5F). The phenotype CD45+TCRb+CD4+, CD45+TCRb+CD8+, and CD45+CD19+, identified CD4, CD8, T cells and B cells respectively, as the population of lymphocytes infiltrating *Pdcd10^BECKO^* brains (Fig 5F). Histological analysis of *Pdcd10^BECKO^* brains showed a significant elevation of IBA1+ microglia cells in the vicinity of CCM lesions (Fig 5G, 5H). In addition, lesion rupture and thrombosis were associated with an increase in IBA1+ microglia cells and IBA1+IB4+ leukocyte infiltration (Fig. 5G, 5H, Supplemental Fig. 5, Supplemental Fig. 7). The IBA1+ microglia cells located nearby the thrombus were hypertrophied and ameboid in morphology and enclosed by GFAP+ astrogliosis (Fig. 5G, 5I). These results confirm that neuroinflammation is a hallmark of mature CCM lesions and that inflammatory chemokines produced by CCM endothelium and A3-activated astrocytes may contribute to attracting innate and adaptive immunity and the propensity to immunothrombosis.

**Fig. 5.**
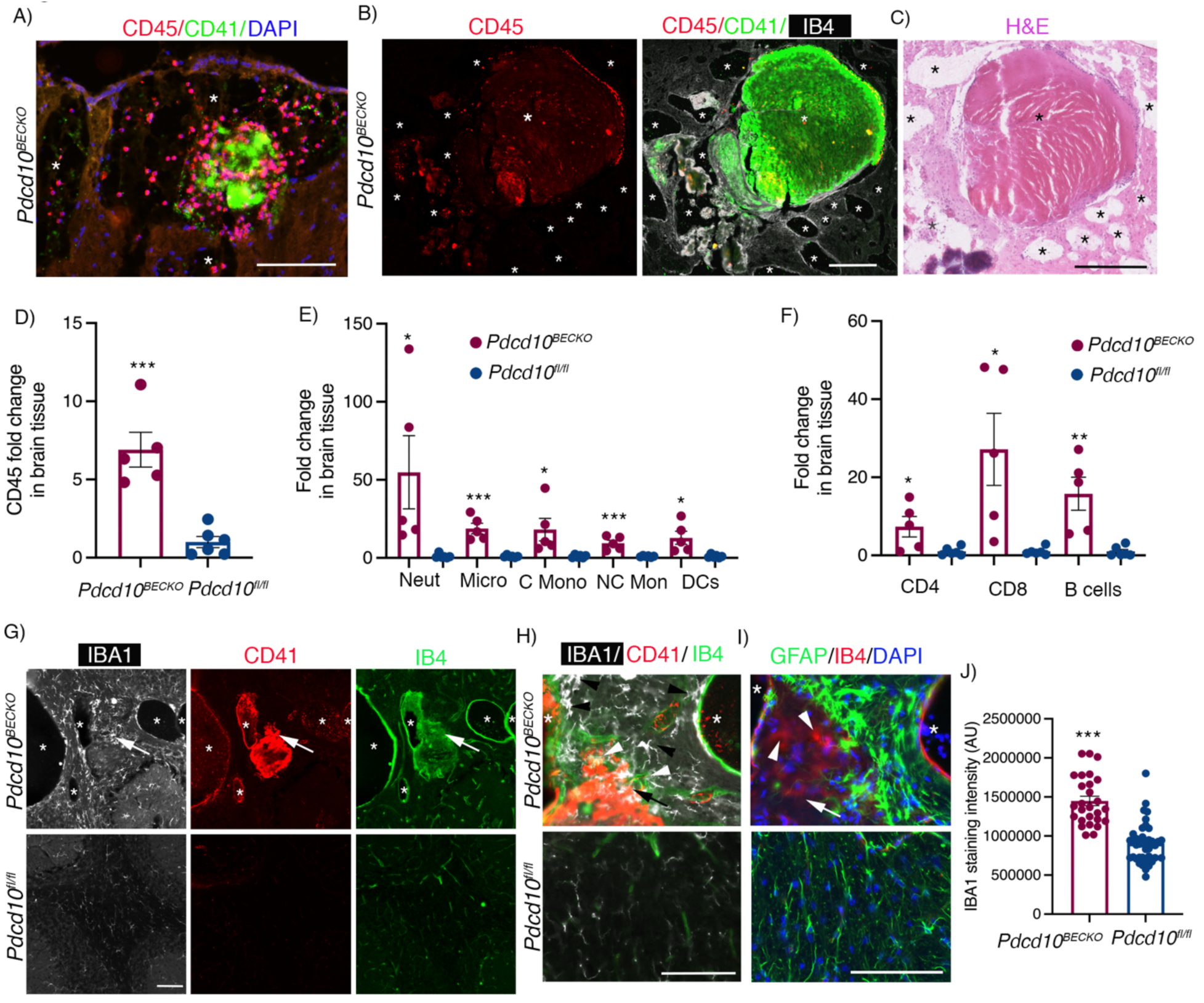
Increased presence of immune cells in CCM brain tissue. **A,** Immunofluorescence analysis shows leukocyte recruitment CD45+ (red) and early platelet aggregation by CD41+ (green) in the vascular lumen of lesions in P75 *Pdcd10^BECKO^* brains. Nuclei are labelled by DAPI (blue). Asterisks denote vascular lumen of CCM lesions. Scale bar, 100 µm. **B,** Large mature vascular thrombosis associated with CD45+ leukocyte infiltration in P75 *Pdcd10^BECKO^* brains. Scale bar, 200 µm. **C,** Hematoxylin and eosin staining of a serial section in *b*. Scale bar, 250 µm. Asterisks denote vascular lumen of CCM lesions. **D,** Flow cytometry of CD45+ cells isolated from P80 *Pdcd10^BECKO^* brains compared to *Pdcd10^fl/fl^* brain controls. **E,** Flow cytometry of myeloid cells isolated from P80 *Pdcd10^BECKO^* brains compared to *Pdcd10^fl/fl^* brain controls. **F,** Flow cytometry of lymphoid cells isolated from P80 *Pdcd10^BECKO^* brains compared to *Pdcd10^fl/fl^* brain controls. Data are mean ± SEM, n = 4 or 5 mice in each group. *, *P*<0.05, **, *P*<0.01, ***, *P*<0.001. **G,** Immunofluorescence analysis for microglia IBA1+ (white), platelets CD41+ (red) cells and labeling of the brain vasculature, using isolectin B4 (green), of cerebral sections from P80 *Pdcd10^BECKO^* brains compared to littermate *Pdcd10^fl/fl^* brain controls. Asterisks denote vascular lumen of CCM lesions. Scale bar, 100 µm. Arrows indicate region at high magnification observed in *h*. **H**, Increase of IBA1+ microglial cells (black arrowheads) and infiltrated IBA1+IB4+ leukocytes (white arrow heads) can be observed near thrombus (CD41+, red) of *Pdcd10^BECKO^* brains. **I,** Immunofluorescence analysis for GFAP (green), and IB4 (red) from a serial section in *g* and *h*, Nuclei are labelled by DAPI (blue). GFAP+ astrocytes enclosed IB4+ cells (white arrowheads) present in the thrombus in *h*. Asterisks denote vascular lumen of CCM lesions. **J,** microglia IBA1+ staining quantification. Scale bar, 100 µm. n = 3 mice in each group.

### *Ikkb^BECKO^* mice show increased thrombosis during CCM disease

The IKK(IkB Kinase)/NF-kB signaling pathway regulates several components of the immune response and inflammation in endothelial cells ^43, 44^, in which endothelial IKKB is the primary kinase mediating NF-kB activation (NF-kB1/p105, NF-kBp65, A20, NEMO) ^45–47^. Since NF-kB inhibition has anti-inflammatory effects *in vivo,* we wanted to investigate whether specific NF-kB inhibition in brain endothelium during CCM disease prevented the neuroinflammation observed in *Pdcd10^BECKO^* brains. Therefore, we produced mice with partial or complete brain endothelial-specific deletion of *Ikkb* (*Ikkb^BECKO/wt^* or *Ikkb^BECKO^*) by crossing mice with the *Ikkb* gene flanked with LoxP sites ^48^ with *Slco1c1-CreERT2;Pdcd10^fl/fl^* mice (*Pdcd10^BECKO^*). Unexpectedly, we noticed that *Pdcd10^BECKO^;Ikkb^BECKO^* mice showed increased mortality approximately at P70 (data not shown). Detailed histological analysis of CCM lesions in stage 1 (single cavern, ∼0.02 mm^2^) and stage 2 (multi cavernous ∼0.12 mm^2^) showed no significant difference in lesion size between brains in *Pdcd10^BECKO^* mice when compared to littermates *Pdcd10^BECKO^;Ikkb^BECKO/wt^* and *Pdcd10^BECKO^;Ikkb^BECKO^* mice (Fig 6A, 6C, 6F, 6H). However, there was a significant increase in the number of stage 1 and stage 2 lesions in *Pdcd10^BECKO^;Ikkb^BECKO/wt^* and *Pdcd10^BECKO^;Ikkb^BECKO^* mice compared to littermates *Pdcd10^BECKO^* mice (Fig 6B, 6D). Moreover, the histological analysis showed an increased presence of thrombosis in stage 2 lesions, but not stage 1 lesion, in *Pdcd10^BECKO^;Ikkb^BECKO/wt^* mice compared to littermate *Pdcd10^BECKO^* mice (Fig. 6E-6H). Brains from *Pdcd10^BECKO^;Ikkb^BECKO^* manifest increased microglia activation and multifaceted CCM lesion formation, which may contribute to the high mortality observed at P70 (Fig. 6I, 6J). These results are consistent with previous findings that indicate that NF-kB inhibition might result in severe proinflammatory effects in particular tissues ^49^, potentially by loss of NF-kB dependent gene expression of genes that participate in the resolution of neuroinflammation ^50–52^, enhance of NLRP3 inflammasome activity ^40^, or by inducing apoptosis ^46^. Further studies will be needed to evaluate pharmacological strategies targeting different nodes of neuroinflammation. Overall, our study suggests that CCM lesions transit through different stages or phases that are histologically and molecularly different and that neuroinflammation plays a critical role in CCM pathogenesis during the chronic stage (Fig. 7). However, therapeutic approaches designed to target brain endothelial NF-kB activity may lead to detrimental side effects.

**Fig. 6.**
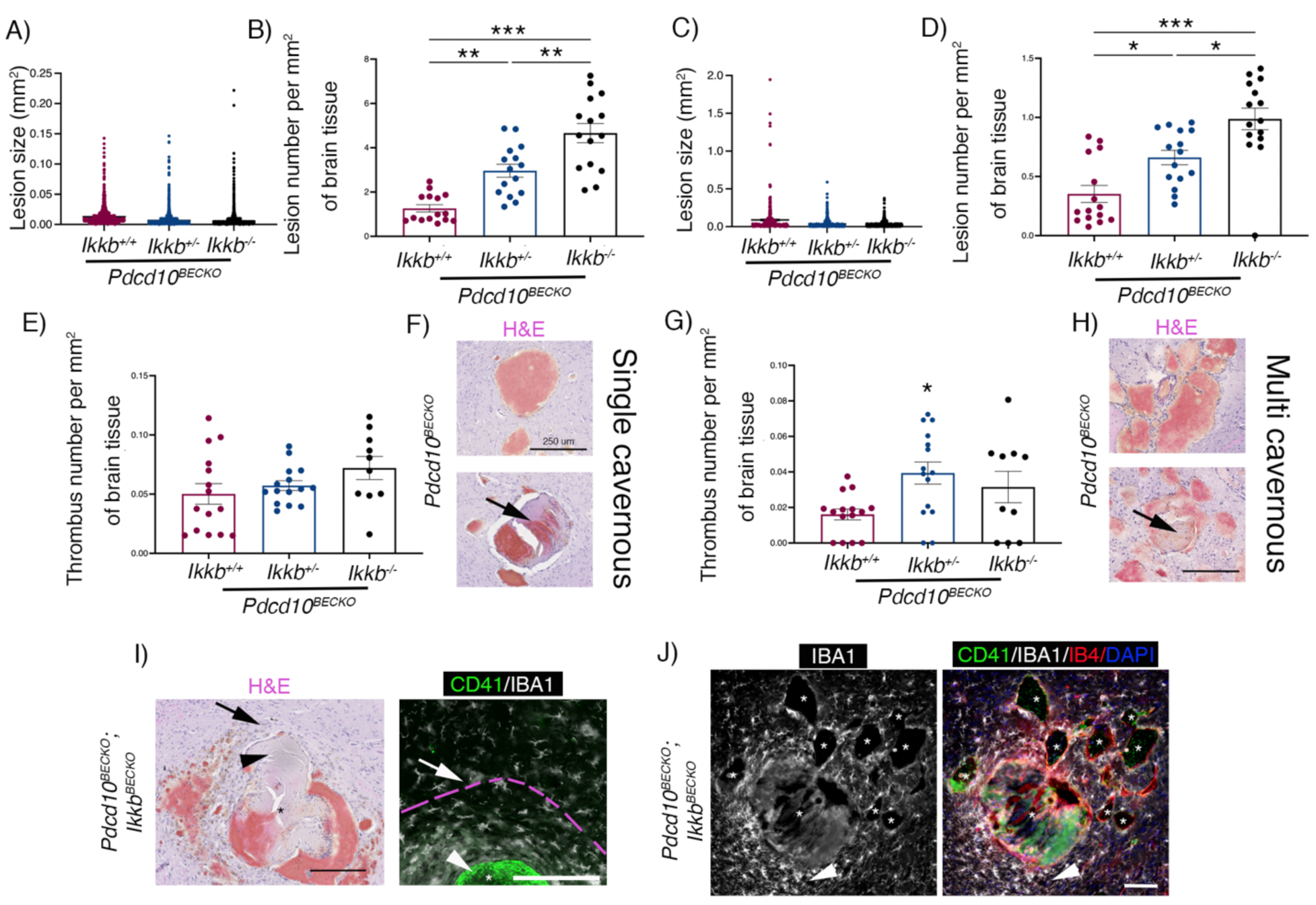
Loss of brain endothelial IKKb increases lesion number and thrombosis during CCM disease. **A,** Analysis and quantification of stage 1 (single cavernous) lesion size in *Pdcd10^BECKO^;Ikkb^+/+^*, *Pdcd10^BECKO^;Ikkb^BECKO/wt^* (*Pdcd10^BECKO^;Ikkb^+/-^*) and *Pdcd10^BECKO^;Ikkb^BECKO^* (*Pdcd10^BECKO^;Ikkb^-/-^*) brains. **B,** Analysis and quantification of the number of stage 1 lesions per mm^2^ in *Pdcd10^BECKO^;Ikkb^+/+^*, *Pdcd10^BECKO^;Ikkb^BECKO/wt^* (*Pdcd10^BECKO^;Ikkb^+/-^*) and *Pdcd10^BECKO^;Ikkb^BECKO^* (*Pdcd10^BECKO^;Ikkb^-/-^*) brains. **C,** Analysis and quantification of stage 2 (multi-cavernous) lesion size in *Pdcd10^BECKO^;Ikkb^+/+^*, *Pdcd10^BECKO^;Ikkb^BECKO/wt^* (*Pdcd10^BECKO^;Ikkb^+/-^*) and *Pdcd10^BECKO^;Ikkb^BECKO^* (*Pdcd10^BECKO^;Ikkb^-/-^*). **D,** Analysis and quantification of the number of stage 2 lesions per mm^2^ in *Pdcd10^BECKO^;Ikkb^+/+^*, *Pdcd10^BECKO^;Ikkb^BECKO/wt^* (*Pdcd10^BECKO^;Ikkb^+/-^*) and *Pdcd10^BECKO^;Ikkb^BECKO^* (*Pdcd10^BECKO^;Ikkb^-/-^*). Animals were injected at P5 with 4-hydroxi-tamoxifen to differentially assess lesion burden^19^. Data are mean ±SEM, n = 3 or 4 mice in each group. **E,** Analysis and quantification of the number of thrombi in stage 1 lesions per mm^2^ in *Pdcd10^BECKO^;Ikkb^+/+^*, *Pdcd10^BECKO^;Ikkb^BECKO/wt^* (*Pdcd10^BECKO^;Ikkb^+/-^*) and *Pdcd10^BECKO^;Ikkb^BECKO^* (*Pdcd10^BECKO^;Ikkb^-/-^*) brains. **F,** Hematoxylin and eosin staining of stage 1 lesions in P80 *Pdcd10^BECKO^* mice. Arrow indicate thrombus present in stage 1 lesion. **G,** Analysis and quantification of the number of thrombi in stage 2 lesions per mm^2^ in *Pdcd10^BECKO^;Ikkb^+/+^*, *Pdcd10^BECKO^;Ikkb^BECKO/wt^* (*Pdcd10^BECKO^;Ikkb^+/-^*) and *Pdcd10^BECKO^;Ikkb^BECKO^* (*Pdcd10^BECKO^;Ikkb^-/-^*). **H,** Hematoxylin and eosin staining of stage 2 lesions in P80 *Pdcd10^BECKO^* mice. The arrow indicate thrombus present in stage 2 lesion. **I,** Hematoxylin and eosin staining show a multifaceted CCM lesion in *Pdcd10^BECKO^;Ikkb^BECKO^* brains, a serial section was used for immunofluorescence analysis for CD41 (green) and IBA1 (white). Arrow indicate similar regions. Increase in IBA1+ microglial cells with hypertrophied and ameboid in morphology are observed to form a ring near the thrombus. Pink broken line indicate area of microglia activation. Arrowheads indicate thrombus. Scale bar, 200 µm. **J,** Immunofluorescence analysis of CD41+ (green), IBA1+ (white) and labeling of the brain vasculature, using isolecting B4 (red), of cerebral sections from P80 *Pdcd10^BECKO^;Ikkb^BECKO^*. Exacerbated increased of IBA1+ microglial cells near a lesion with thrombosis (arrowhead). Lesions with not thrombosis show a moderate recruitment of microglial cells. Nuclei are labelled by DAPI (blue). Asterisks denote vascular lumen of CCM lesions. Animals were injected at P1 with 4-hydroxi-tamoxifen. Scale bar, 100 µm. Data are mean ±SEM, n = 3 or 2 mice in each group. *, *P*<0.05, **, *P*<0.01, ***, *P*<0.001.

**Fig. 7.**
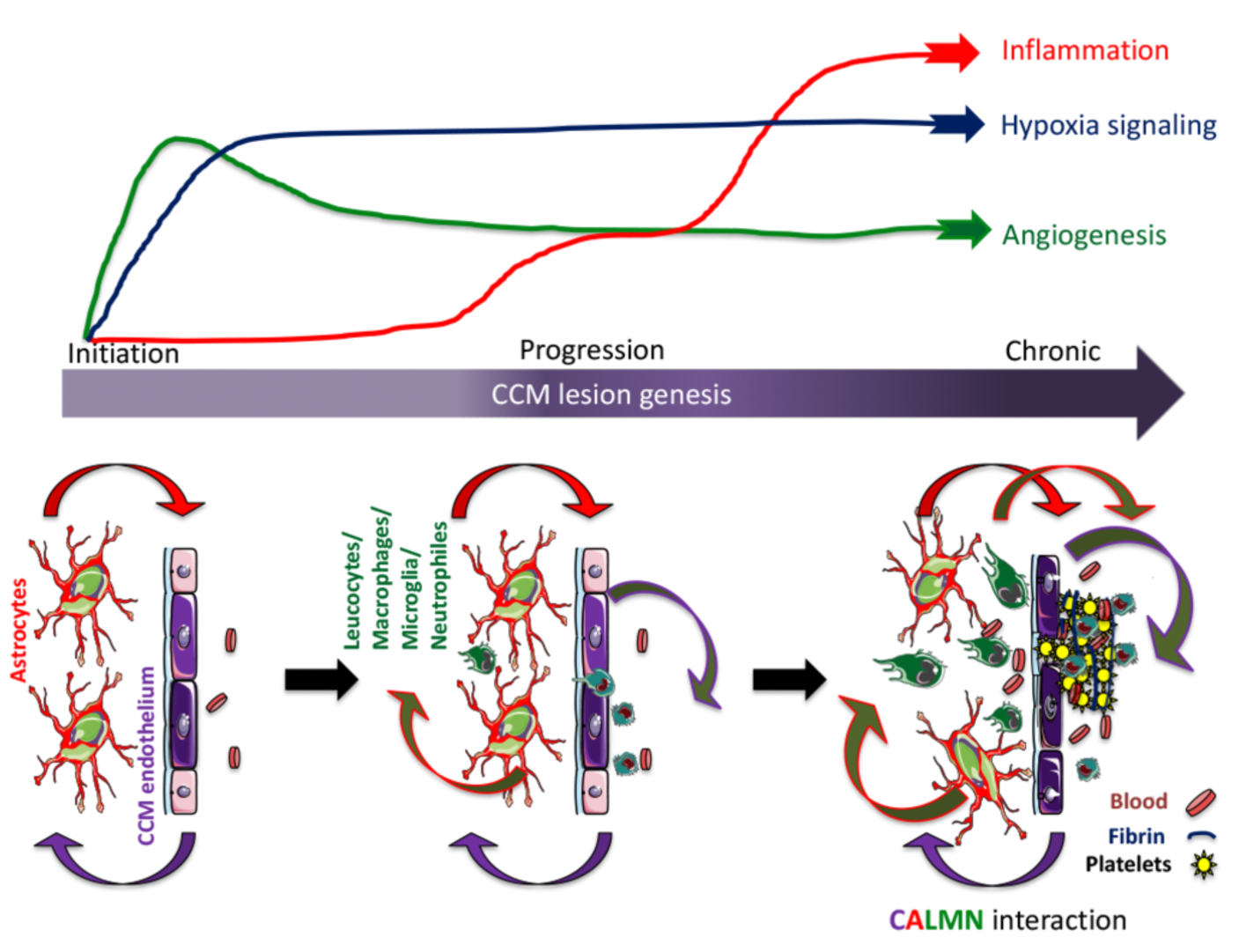
CALMN interaction plays a critical role in the pathogenesis of CCM disease. The illustration shows that angiogenesis and hypoxia signaling are essential in CCM lesion formation (initiation)^22^, while the inflammation pathway initiates in the progression phase and may contribute to mature active CCM lesions, lesion growth, immunothrombosis, and bleedings. Our model proposes that CCM endothelium and astrocytes synergize to recruit inflammatory cells to CCM lesions. Moreover, a reciprocal interaction between CCM endothelium, astrocytes, leukocyte, and macrophage/microglia, neutrophils, that we termed CALMN interaction, is critical for the transition of lesions into aggravating active lesions.

## Discussion

We show here that neuroinflammatory signaling pathways from astrogliosis and CCM endothelium are contributors to immune cell recruitment and thrombi formation in vascular lesions of a mouse model of CCM disease (Fig. 7). Through RNA-seq analysis from astrocyte-translated mRNA, we identified that astrocytes in the presence of CCM lesions acquire a reactive phenotype of neuroinflammatory astrogliosis involved in the production of leukocyte chemotactic factors, antigen presentation, and inflammatory response. We show that CCM endothelium plays a crucial role in recruiting inflammatory cells and promoting thrombi formation potentially through chemotaxis and NLRP3 inflammasome activity. Additionally, our analysis reveals that neuroinflammation conferred by neuroinflammatory astrocytes and CCM endothelium attracts innate and adaptive immunity during CCM disease. Our study proposes that while hypoxia and angiogenesis pathways are essential in CCM lesion formation, inflammation and NLRP3 inflammasome activity play a more critical role in transitioning into mature and active CCM lesions. Our findings introduce the concept that “CALMN (CCM endothelium, astrocytes, leukocytes, monocytes/microglia, neutrophils) interaction” contribute to the pathogenesis of CCMs and a better understanding of the CALMN interaction may lead to new therapeutic strategies for CCMs disease (Fig. 7).

Our study reveals previously unknown links between neuroinflammatory astrocytes and inflamed CCM endothelium as contributors that trigger leukocyte recruitment and facilitates immunothrombosis in CCM lesions. Our work and others have shown that angiogenesis signaling pathways are crucial in CCM lesion formation characterized by isolated single caverns that range in size (stage 1) ^19–22, 53^. Astrocytes, the most abundant cell type in the CNS, respond to the CCM endothelium and propel CCM formation in a non-cell-autonomous manner, mediated by hypoxia and angiogenesis programs ^22^. However, the mechanistic linkage by which CCM lesions acquire the multicavernous stage (stage 2), characterized by hemosiderin deposits and immune cell infiltration in CCM brains of both humans and mice ^17^ remains unclear. Studies have shown that polymorphisms in inflammatory pathways (e.g., Tlr4 signaling) contribute to variability in CCM disease severity ^54–56^ and the depletion of B lymphocytes in a genetically sensitized CCM mouse model reduces the maturation and propensity for chronic bleedings ^57^. Recent studies in human and mouse models show that inflammation contributes to brain vascular malformations ^22, 23, 57–59^. These observations provide direct evidence that inflammation plays a critical role in CCM lesion maturation into the clinical manifestation of CCM disease. Our findings have important implications for explaining the mechanism behind CCM lesions maturation, by demonstrating that, in addition to CCM hemorrhage, the neuroinflammatory signaling pathway triggered by both astrocytes and CCM endothelial cells significantly contributes to lesion progression, during which inflammatory cell infiltration and thrombosis are observed. Indeed, an increase in astrocyte CX3CL1 signaling may play an important role in recruiting both microglia and peripheral inflammatory cells into CCM lesions via the receptor CX3CR1^60^. Moreover, the increase of astrocyte MCP1 (*Ccl2*), a chemoattractant for monocytes and activated T cells, may synergize with CX3CL1 to recruit the innate and adaptive immune response^58, 61^ as well as to induce disassembly of tight junctions in brain endothelial cells during CCM disease^8, 62^. We observed that neuroinflammatory astrocytes induced in CCMs expressed high mRNA levels of CD74, a cell-surface form of the MHC class-II invariant chain. CD74 expression in astrocytes regulates the expression of COX-2 (*Ptgs2*)^63^, an angiogenic and pro-inflammatory enzyme, which inhibition ameliorates CCM disease ^22^. Astrocyte reactivity in diseases that affect the CNS is well documented ^64^. Indeed, astrocytes are proposed to exist in two distinct reactive states, such as neuroinflammatory-A1 or neuroprotective-A2 reactive astrocytes. However, more than two populations of reactive astrocytes may also coexist ^32–34^. Recent studies using single-cell analysis and spatial transcriptomics have demonstrated the significant heterogeneity of astrocytes in the brains of mice following systemic injection with endotoxin lipopolysaccharide (LPS) ^65^. Our data showed that, during CCM disease, there is no clear polarization of astrocytes toward specific A1- like or A2-like reactive states (Supplemental Fig. 3). Instead, we observed that astrocytes activated during vascular disease acquire a particular transcriptional signature that we denominated A3 (n+1 astrocyte,^33^) CCM reactive astrocyte with neuroinflammatory capacity. These findings suggest that astrocyte polarization induced by CCM endothelium differs from the one induced by LPS and pro-inflammatory cytokines ^32^. This work proposes that future studies should aim to identify a selective mechanism of A3 reactive astrocytes inhibition in the context of CCM, which may constitute a novel therapeutic target for the disease ^22^. In addition, we identified that A3 reactive astrocytes express high levels of genes directly associated with neurodegenerative diseases with cerebral angiopathies, such as *Ndufa4l2*, *App*, *Cx3cr1, S1pr3* ^66, 67^. The manipulation and further characterization of the role of A3 reactive astrocytes in CCMs and other cerebrovascular malformations will be an important area for future research.

In CCMs, brain endothelial cells show gross endothelial changes (CCM endothelium) ^5, 6, 8, 10^, leading to blood-brain barrier dysfunction that contributes to neurological deficits and, in some cases, death ^68–70^. Moreover, previous work has demonstrated that, during lesion formation, the CCM endothelium is hypersensitive to angiogenesis and can induce a hypoxic program associated with changes in angiogenesis, inflammation, and endothelial-cell metabolism under normoxic conditions ^22^. Interestingly, our data regarding CCM endothelium at the chronic stage of the disease shows enrichment in genes relevant to mature and active CCM lesions. These changes included the elevation of neutrophil degranulation, platelet activation, ROS production^71^, and NLRP3 inflammasome. In addition, CCM endothelium in the chronic stage shows significant upregulation of transcripts coding for chemokines such as MCP1, CCL5, CX3CL1, known for recruiting peripheral leukocytes into inflamed tissue ^61, 72^. We also found that genes implicated in hypoxia signaling such as APLN, EGLN3, VEGFa, LOXL2, CD44 were highly increased in chronic CCM endothelium and may participate in disease severity. The data establish that the potential mechanistic crosstalk between astrocytes and CCM endothelium can trigger recruitment of inflammatory cells, arising from both brain parenchyma (microglia) and periphery immune system (leukocytes) into mature CCM lesions, that could potentially propagate lesion growth, thrombosis, and bleeding. Accordingly, altered levels of several genes associated with blood coagulation, TSP1 (*Thbs1*), EPCR (*Procr*), TM (*Thbd*), VWF, tPA (*Plat*), indicate that in the chronic model, some CCM lesions may have a propensity to bleeding (high levels in EPCR and TM and TFPI) while others may have a propensity to become thrombotic (high levels in PAI-1, TSP1, VWF, low in tPA, and inflammation) ^23, 73^. It is also possible that pro-thrombotic signals in chronic CCM lesions overcome the local anti-coagulant vascular domain ^25^ and promote immunothrombosis in largely inflamed lesions. Notably, these findings in our CCM mouse model are consistent with human studies in which inflammatory, anti-coagulant, and angiogenic plasma soluble proteins were shown to be promising predictive and prognostic biomarkers during CCM disease progression ^1, 25, 74^.

In our study, transcriptomic data from CCM endothelium in the chronic stage of the disease suggest a critical role for genes associated with vascular homeostasis, aneurysms, neurodegenerative diseases, and senescence genes, including IL6, CD74, COL10A1, SERPINA1e/b/d, CYP3A13, GKN3, SERPINE1. Future studies could assess the impact of these genes in CCM lesion genesis and their contribution to neurological manifestation of the disease in animal models. In addition, our histological and gene expression analysis suggest a critical role for the endothelial NLRP3 inflammasome activity, which may significantly contribute to both the inflammation and thrombosis ^75^ observed in CCM chronic lesions. Moreover, persistent inflammasome activation in endothelium and immune cells contributes to neurological and neurodegenerative conditions ^76^ that could be relevant to the clinical symptomatology of CCM patients. Therefore, future research should focus on further identifying the molecular mechanisms that contribute to the increase of NLRP3 during CCM disease and investigate the therapeutic effect of drugs targeting NLRP3 inflammasome signaling ^77^ in CCM disease. Indeed statins, aspirin and inhibitors of ROS activity, negatively regulate NLRP3 inflammasome activity^77, 78^ and have been identified as a potential therapy for CCM disease in pre-clinical and clinical studies^79–81^. It is possible that different mutations in CCM genes (CCM1, CCM2 or CCM3) may result in different biological mechanisms that lead to lesion aggressiveness. However, the current evidence suggests that mutation of any CCM genes operates on similar pathways during disease initiation^1, 17, 22, 55, 57, 82^. Consistently, CCM in human and murine studies has shown that neuroinflammation contributes to pathogenesis in CCM1 (most common) and CCM3 (most aggressive) diseases ^17, 54, 57, 83, 84^. Lastly, in addition to identifying potential therapeutic strategies to ameliorate CALMN interaction and inflammatory responses during the chronic stage of CCM disease, we propose that therapeutic approaches targeting brain endothelial NF-kB activity during CCM might contribute to detrimental side effects.

---

**Supplemental Fig. 1.**
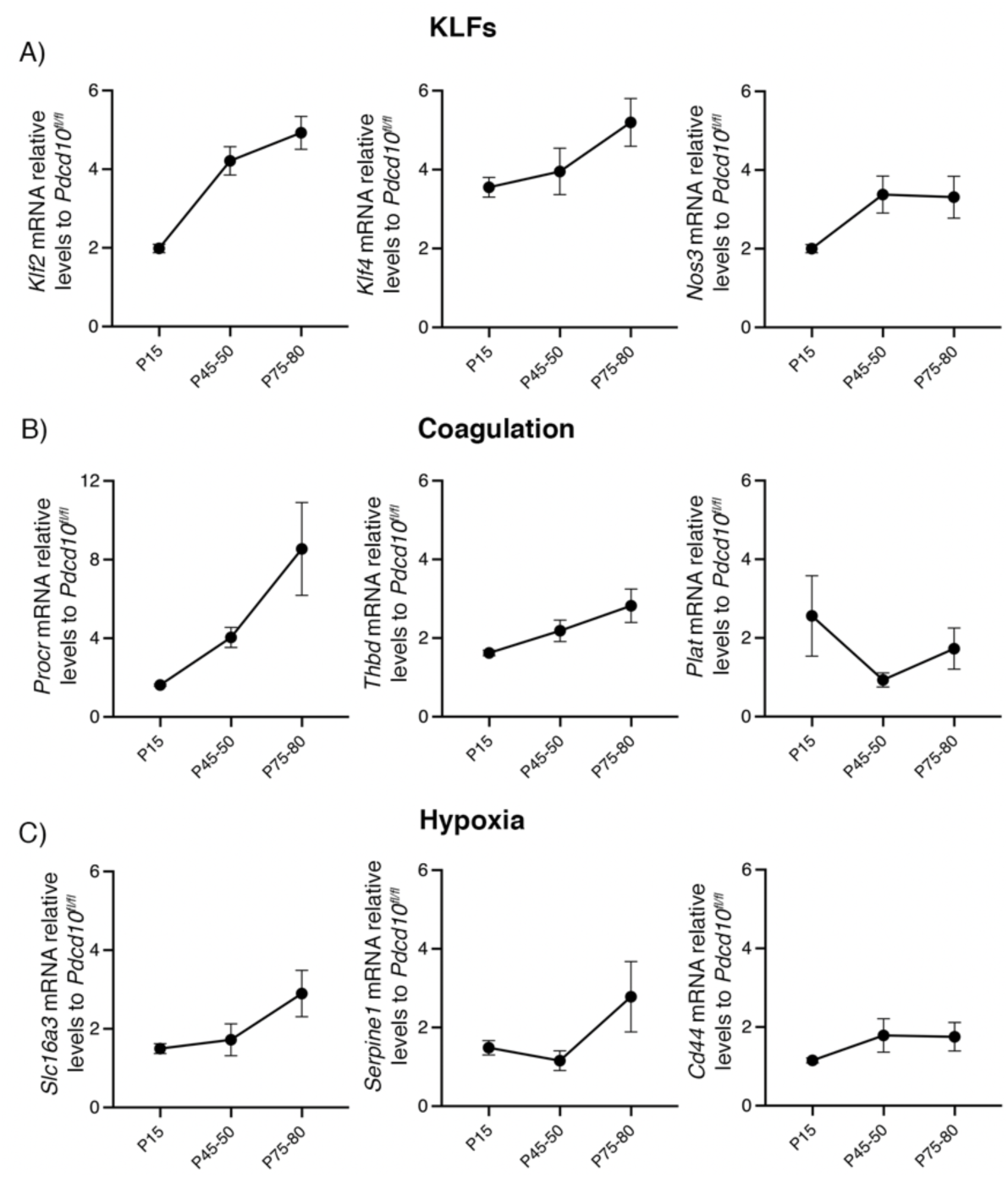
Increase of hypoxia and coagulation signaling pathways in CCM disease. **A,** CCM lesions are present in the cerebellum and cerebrum of *Pdcd10^BECKO^* mice at acute (P15), progression (P50), and chronic stage (P80). **B,** Analysis of *Klf2*, *Klf4*, *Nos3*, *Procr*, *Thbd*, *Plat*, *Slc16a3, Serpine1,* and *Cd44* mRNA levels by RT-qPCR in cerebral tissue from mice at chronic stage in samples from Fig 1b. Data are mean ±SEM, n = 3 or 6 mice in each group.

**Supplemental Fig. 2.**
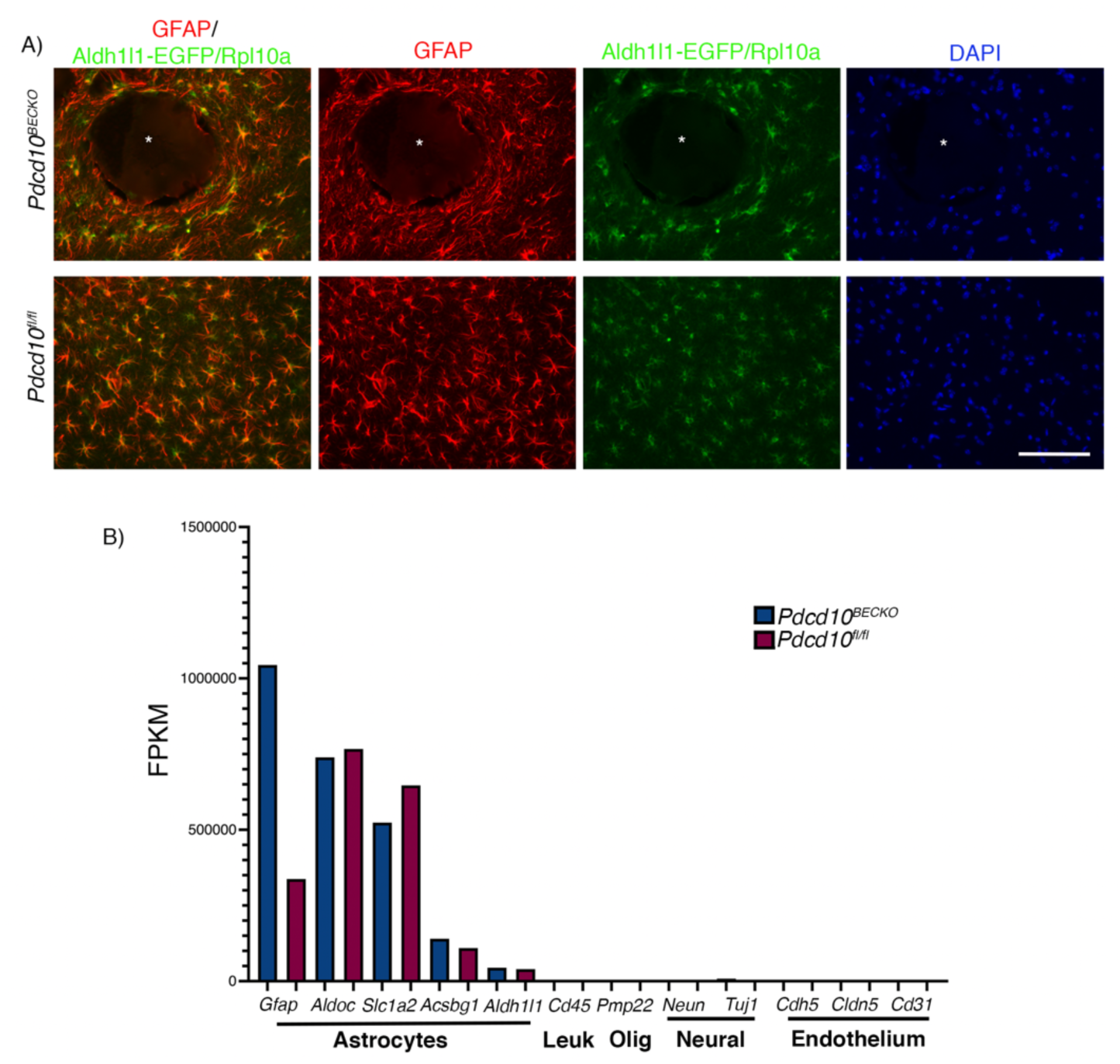
Neuroinflammatory astrocytes in CCM disease. **A,** Immunofluorescence analysis show colocalization of GFAP+ astrocytes with cells expressing EGFP-RpL10a in *Pdcd10^fl/fl^;Aldh1l1-EGFP/Rpl10a* brains sections. Asterisks denote the vascular lumen of CCM lesions. Scale bar is 100 µm. n = 3 mice in each group. **B,** Analysis of known brain cell-specific gene markers from *Pdcd10^BECKO^;Aldh1l1-EGFP/Rpl10a* (blue bar) and *Pdcd10^fl/fl^;Aldh1l1-EGFP/Rpl10a* littermate controls (red bar) following ribosome-bound mRNA analysis by RNA-seq. n = 3 in each group.

**Supplemental Fig. 3.**
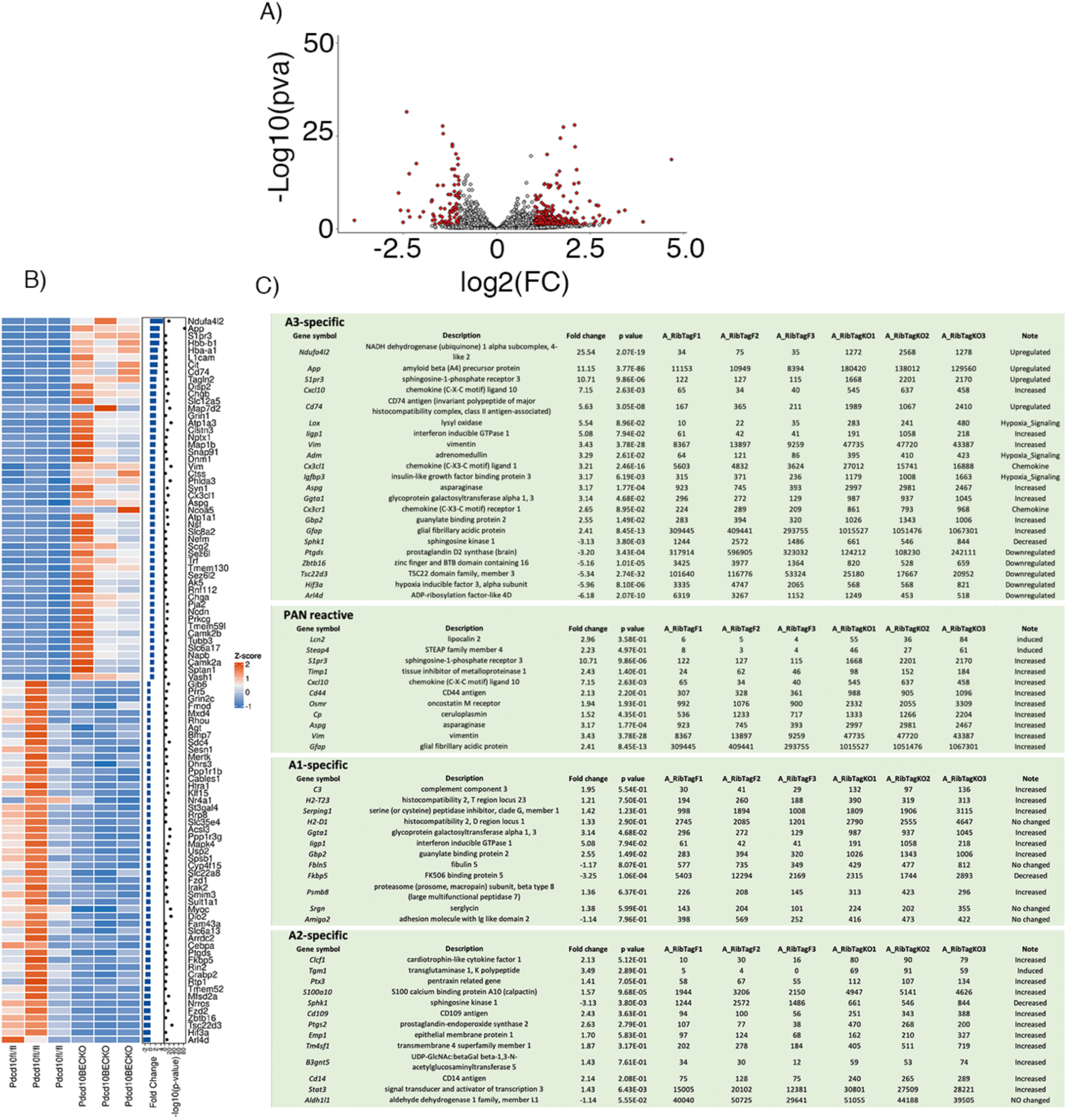
A3 astrocytes in CCM disease. **A,** Volcano plot of differentially expressed transcripts in P75 *Pdcd10^BECKO^;Aldh1l1-EGFP/Rpl10a* versus littermate control *Pdcd10^fl/fl^;Aldh1l1-EGFP/Rpl10a.* Transcripts are represented on a log2 scale in Fig 2B. The significantly down- and up-regulated genes are labeled in red. n = 3 mice in each group. **B,** List of the top 50 up- or down-regulated genes from translated mRNAs obtained from ribosomes in astrocytes from *Pdcd10^BECKO^;Aldh1l1-EGFP/Rpl10a* brains in Fig 2D. Fold change and *P* values are shown for each gene. Heatmaps comparing the mean expression of DEG. n = 3 mice in each group. **C,** Transcriptional signature of neuroinflammatory 3A astrocytes in CCM disease. Pan reactive, A1 specific and A2 specific genes are shown. Fold change and *P* values are shown for each gene and reads values for each biological replicate. n = 3 mice in each group.

**Supplemental Fig. 4.**
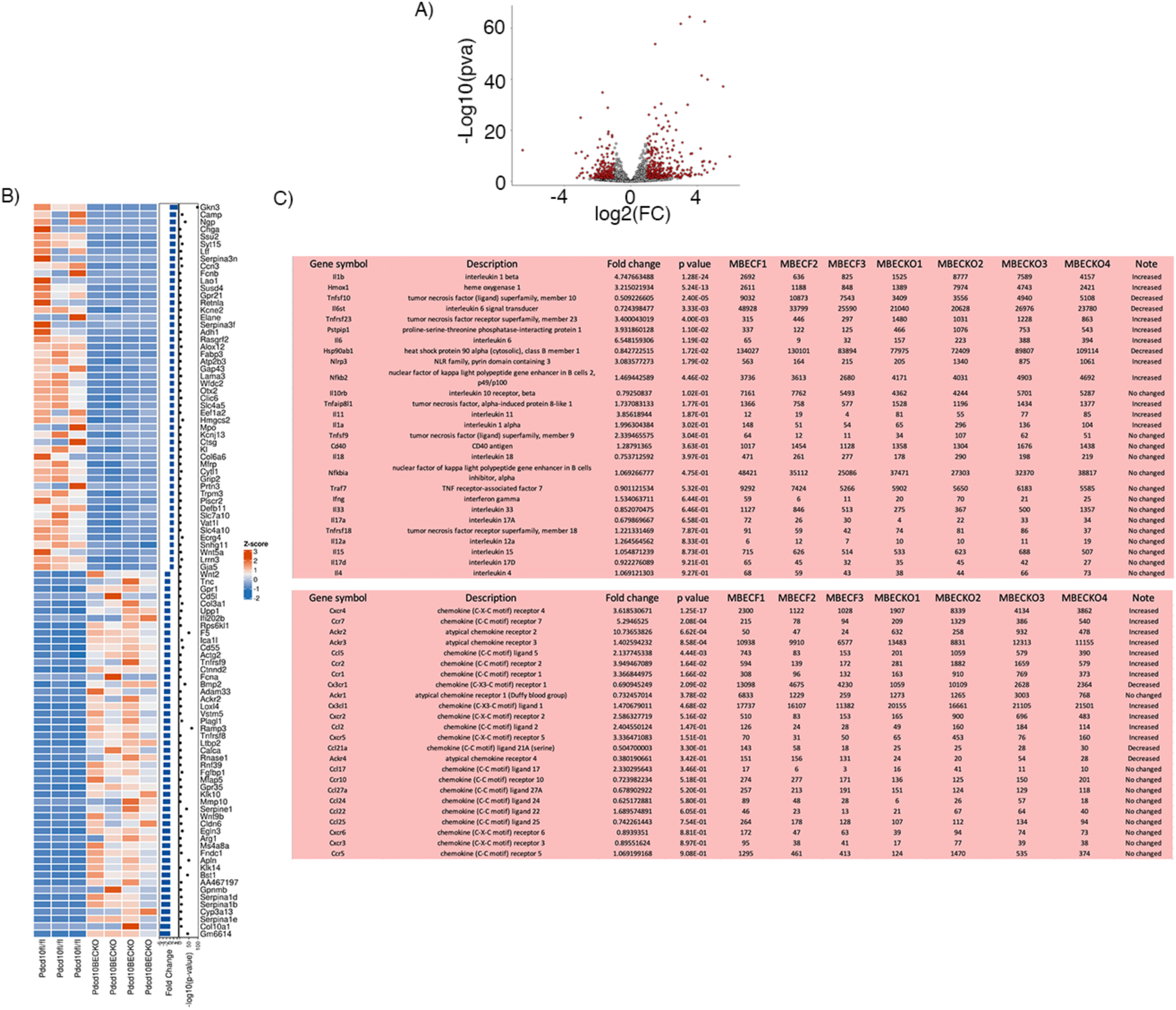
Inflammation pathway is increased in the CCM endothelium. **A,** Volcano plot of differentially expressed transcripts in fresh isolated brain endothelial cells from P75 *Pdcd10^BECKO^;Aldh1l1-EGFP/Rpl10a* versus isolated brain endothelial cells from littermate control *Pdcd10^fl/fl^;Aldh1l1-EGFP/Rpl10a.* Transcripts are represented on a log2 scale from Fig. 3A. **B,** List of the top 50 up- or down-regulated genes from isolated brain endothelial cells obtained from *Pdcd10^BECKO^;Aldh1l1-EGFP/Rpl10a* brains. Fold change and *P* values are shown for each gene. n = 3 or 4 mice in each group. Heatmaps comparing the mean expression of DEG. **C,** Analysis of inflammatory signaling from *Pdcd10^BECKO^;Aldh1l1-EGFP/Rpl10a* isolated brain endothelial cells. Fold change and *P* values are shown for each gene. n = 3 or 4 mice in each group.

**Supplemental Fig. 5.**
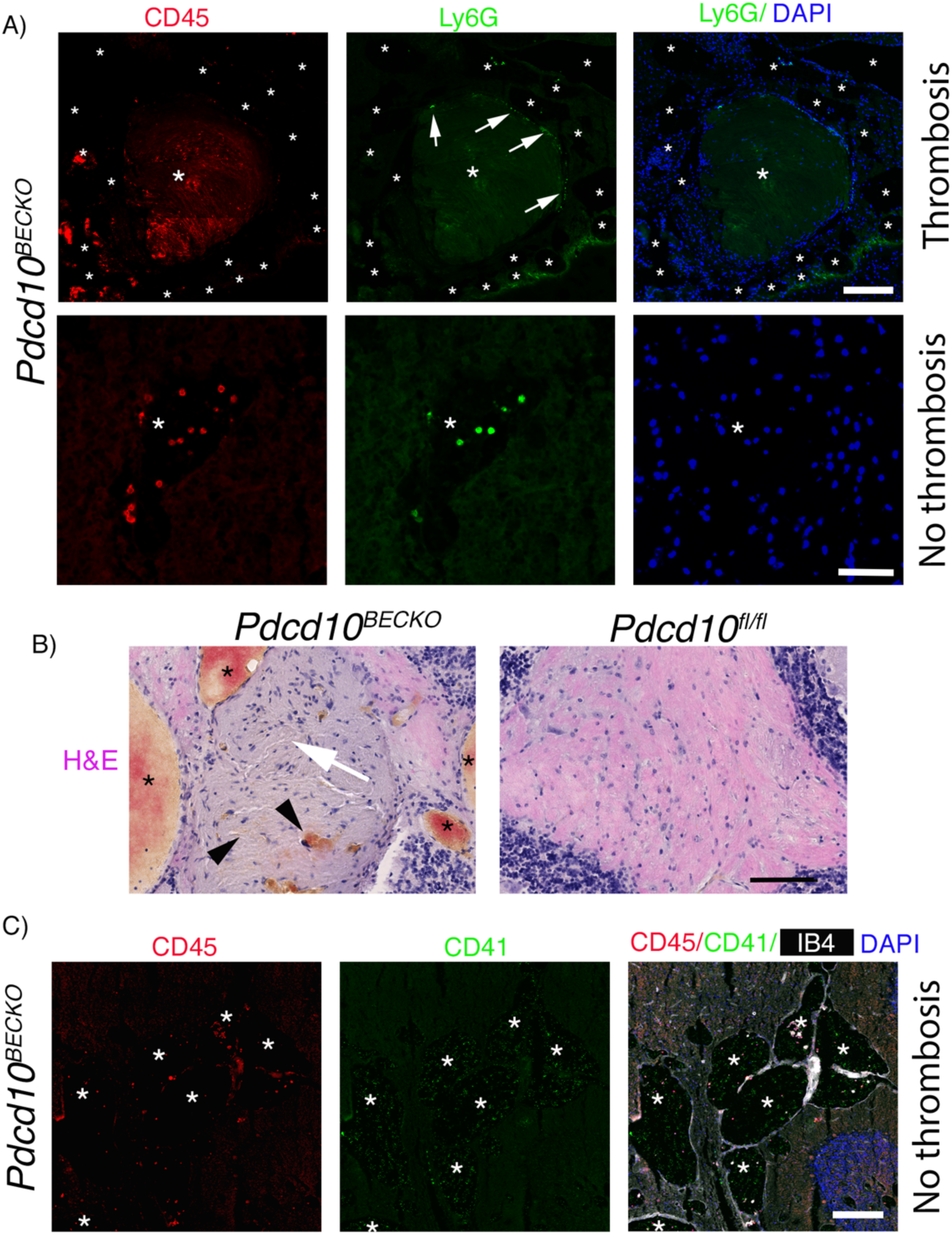
Increased presence of CD45+ and Ly6G+ cells in CCM brain tissue. **A,** Immunofluorescence analysis from a serial section used in Fig. 5b and 5c shows leukocyte recruitment CD45+ (red) and accumulation of Ly6G+ neutrophils (green, arrows) in the vascular lumen and vessel wall of lesions in P75 *Pdcd10^BECKO^* brains, respectively. Non-thrombus lesions also show the presence of Ly6G+ neutrophils (green). Nuclei are labelled by DAPI (blue). Asterisks denote vascular lumen of CCM lesions. Scale bar, 200 µm (top) and 50 µm (bottom). **B,** Hematoxylin & eosin staining of a serial section in Fig. 5g, Arrow indicates an area of thrombosis, arrowheads indicate bleeding, and asterisks denote the vascular lumen of CCM lesions. Scale bar is 100 µm. **C,** Immunofluorescence analysis for leukocytes CD45+ (red), platelets CD41+ (green) cells and labeling of the brain vasculature, using isolecting B4 (white), of non-thrombotic multi-cavernous lesion in *Pdcd10^BECKO^* brains. Asterisks denote vascular lumen of CCM lesions. Scale bar, 100 µm.

**Supplemental Fig. 6.**
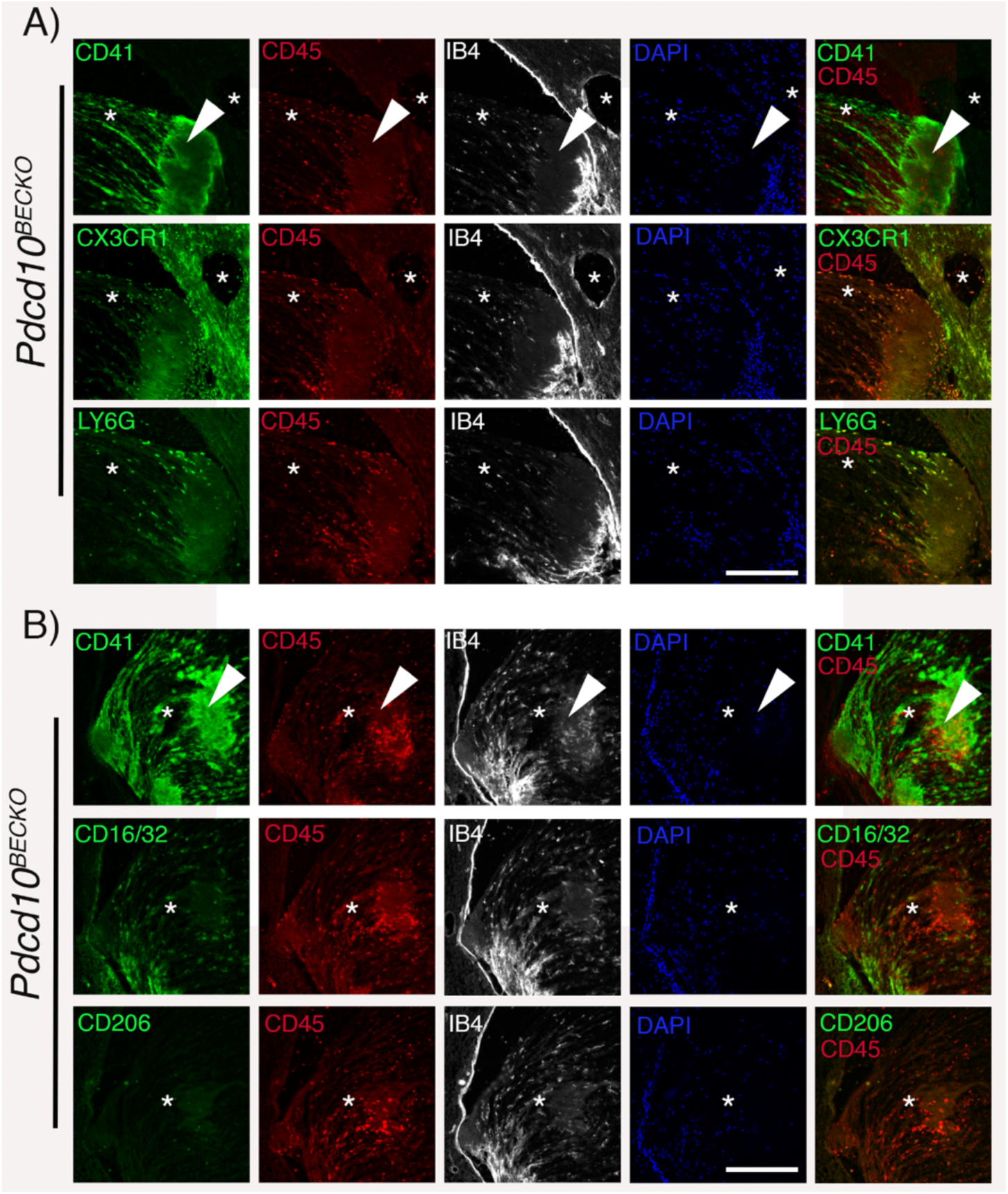
Presence of CD45+, CX3CR1+, Ly6G+, and CD16/32+ cells in the thrombosis area of CCM brain tissue. **A,** Immunofluorescence analysis from serial sections shows leukocyte recruitment CD45+ (red) and accumulation of CX3CR1+ monocytes, Ly6G+ neutrophils in the thrombus (CD41+, green) formed in the vascular lumen of lesions in P75 *Pdcd10^BECKO^* brains. Asterisks denote vascular lumen of CCM lesions. Arrowhead denote vascular thrombus. Scale bar, 200 µm. Nuclei are labeled by DAPI (blue). **B,** Immunofluorescence analysis from serial sections shows leukocyte recruitment CD45+ (red) and accumulation of CD16/32+ monocytes, but not CD206+ monocytes in the thrombus (CD41+, green) formed in the vascular lumen of a lesion in P75 *Pdcd10^BECKO^* brains. Asterisks denote vascular lumen of CCM lesions. Arrowhead denote vascular thrombus. Scale bar, 200 µm. Nuclei are labeled by DAPI (blue).

**Supplemental Fig. 7.**
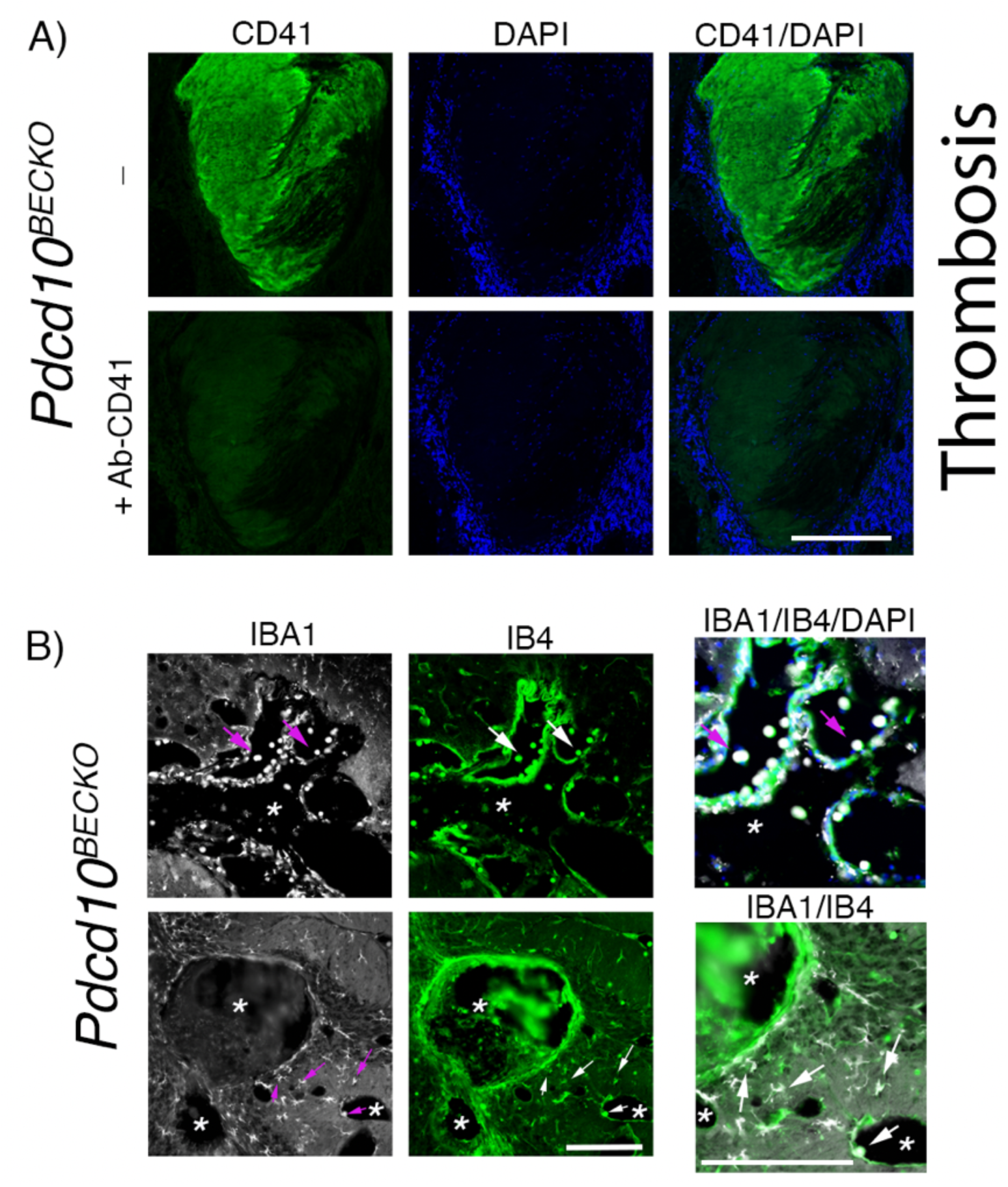
Immunofluorescence of CD41+ thrombus and IBA1+IB4 leukocytes in the vascular lumen and brain parenchyma of CCM brain tissue. **A,** Immunofluorescence analysis of a thrombus (CD41+, green) formed in the vascular lumen of a lesion in P80 *Pdcd10^BECKO^* brain. To confirm CD41+ immunoreactivity, a serial section was pre-treated with a monoclonal anti-CD41 antibody non-labeled (MWReg30) or vehicle for 1 h at room temperature. After washes, the brain tissue was incubated with a CD41-Alexa-488 antibody (MWReg30). Asterisks denote vascular lumen of CCM lesions. Nuclei are labeled by DAPI (blue). Scale bar, 200 µm. **B,** Immunofluorescence staining of IBA1+ microglia and myeloid cells (white), and endothelial marker isolectin B4 (IB4; green) of cerebral sections from P80 *Pdcd10^BECKO^*. Immunofluorescence analysis shows IBA1+IB4+ leukocytes (Arrows) in the lumen and brain parenchyma in P80 *Pdcd10^BECKO^* brains. Asterisks denote vascular lumen of CCM lesions. Scale bar, 200 µm.

## Material and methods

### Genetically modified animals

Brain endothelial-specific conditional *Pdcd10*-null mice were generated by crossing a *Slco1c1* promoter-driven tamoxifen-regulated Cre recombinase (*Slco1c1-CreERT2,* a gift from Markus Schwaninger, University of Lübeck) strain with loxP-flanked *Pdcd10* (*Pdcd10^fl/fl^*, a gift from Wang Min, Yale University;*Slco1c1-CreERT2;Pdcd10^fl/fl^*) mice. Brain endothelial-specific conditional *Ikkb*-null mice were generated by *Slco1c1-CreERT2* strain with loxP-flanked *IKKb* (*Ikkb^fl/fl^*, a gift from Michael Karin, UCSD;*Slco1c1-CreERT2;Ikkb^fl/fl^*). On a postnatal day 1 (P1), mice were administered 50 μg of 4-hydroxy-tamoxifen (H7904, Sigma-Aldrich) by intragastric injection to induce genetic inactivation of the endothelial *Pdcd10* gene in littermates with *Slco1c1-CreERT2;Pdcd10^fl/fl^* (*Pdcd10^BECKO^*), and *Pdcd10^fl/fl^* mice were used as littermate controls. Injection of 4-hydroxy-tamoxifen at P5 was also performed and led to reduced CCM burden^19^ and was used to assess lesions when the brain endothelial *Ikkb* gene was inactivated. We also used non-injected *Slco1c1-CreERT2;Pdcd10^fl/fl^* mice as littermate controls whose gene expression and histology were the same as in *Pdcd10^fl/fl^* mice. To perform Translational Ribosome Affinity Purification (TRAP) in astrocytes ^27, 28^, we generated *Slco1c1-iCreERT2;Pdcd10^fl/fl^;Aldh1l1-EGFP/Rpl10a* and controls littermate *Pdcd10^fl/fl^;Aldh1l1-EGFP/Rpl10a.* All animal experiments were performed in compliance with animal procedure protocols approved by the University of California, San Diego Institutional Animal Care and Use Committee.

### Astrocyte ribosome isolation

*Aldh1l1-EGFP/Rpl10a* mice crossed with the CCM mouse model were used to isolate TRAP mRNAs ^27, 28^ from astrocytes using the protocol and instructions as previously described ^85^. Astrocyte-TRAP mRNAs were from brains of mice age P75.

### Brain endothelial cell (BEC) isolation

Adult P75 *Pdcd10^BECKO;^Aldh1l1-EGFP/Rpl10a* mice (*Pdcd10^BECKO^*) and *Pdcd10^fl/fl^*;*Aldh1l1-EGFP/Rpl10a* (*Pdcd10^fl/fl^*) control littermates were sacrificed, and their brains were isolated and placed into cold solution A (0.5% bovine serum albumin (BSA) in DMEM and 1 μg/μl glucose, 10mM HEPES, 1x penicillin-streptomycin). Meninges and choroid plexus were removed, and one brain of *Pdcd10^BECKO^* mice was minced using scissors in cold solution A. We used two brains of *Pdcd10^fl/fl^* mice that were pooled together to collect enough microvasculatures. Brain tissue suspension was then centrifuged at 1000g for 5 minutes at 4oC. The supernatant was removed and the tissue was digested with a collagenase/dispase solution (1mg/ml collagenase/dispase [Sigma-Aldrich], 20 units/ml DNase I [Sigma-Aldrich], and 0.150μg/ml tosyl-lysine-chloromethyl-ketone [Sigma-Aldrich] in DMEM])(51) at 37°C for 1 hour with vigorous shaking every 10 minutes. Tissue suspension was triturated using thin-tipped Pasteur pipettes until fully homogenous and centrifuged at 700g for 5 minutes at 4°C. The supernatant was removed, and the pellet was resuspended in 20ml of 25% BSA solution followed by centrifugation at 2000g for 20 minutes at 4°C. Capillary fragments were pulled down to the bottom of the tube, remaining BSA and myelin were discarded, and the pellet was resuspended in cold solution A followed by centrifugation at 700g for 5 minutes at 4°C. The supernatant was removed, and capillary fragments were incubated in collagenase/dispase solution at 37°C for 1 hour. Solution A was added to inactivate enzymatic activity, and the suspension was centrifuged once at 700g for 5 minutes at 4°C. The cell pellet was resuspended in ACK lysis (Lonza) buffer to lyse red blood cells, and then cells were centrifuged once at 700g for 5 minutes at 4°C. The supernatant was removed, and cells were then incubated with anti-CD45 coated beads and passed through a column, following the manufacturer’s protocol (Miltenyi Biotec). Isolated BECs were recovered by negative selection and used for RNA-seq and histology analysis.

### Leukocyte isolation and flow cytometry analysis

*Pdcd10^BECKO^* mice and *Pdcd10^fl/fl^* control littermates were sacrificed, brains were collected and placed into cold solution A. Meninges, brain stem, white matter/brainstem were removed. Brain tissue was triturated using scissors and centrifuged at 1000g for 5 minutes. The supernatant was removed, and the pellet was incubated in pre-warmed collagenase/dispase solution for 40 minutes at 37°C (with vigorous shaking every 10 minutes). After incubation, cold solution A was used to stop enzymatic activity, and the suspension was centrifuged at 600g for 10 minutes at 4°C. The supernatant was removed, and the cell pellet was resuspended into a Percoll gradient (70% Percoll, 37% Red Percoll, 30% Percoll, respectively) as previously described by Guldner et al. ^86^, followed by centrifugation at 1000g for 25 minutes at room temperature. The final solution consisted of three distinct layers, with a buffy leukocyte layer at the interface of clear 70% Percoll and 37% Red Percoll and a thick myelin layer at the top of the tube. The leukocyte layer was extracted and resuspended in cold 1X HBSS in a fresh tube, followed by centrifugation at 600g for 7 minutes at 4°C. The supernatant was removed, and cells were resuspended in flow cytometry buffer (0.5% BSA, 2mM EDTA in PBS1X).

### Flow cytometry characterization of immune cells

After isolation cells were stained in the dark for 30 min at 4C with a mixture of anti-mouse CD45-PerCP (clone 30-F11, Biolegend), CD11b-PE-Cy7 (clone M1/70, Biolegend), TCRβ-PE-Alexa Fluor610 (clone H57-597, Biolegend), CD4-BV786 (clone GK1.5, Invitrogen), CD8-BV421 (clone 53-6.7, Biolegend). CD19-APC-Cy7 (clone 6D5, Biolegend), Ly6G-FITC (clone 1A8, BD Biosciences), F4/80-BV711 (clone BM8, Biolegend), CD206-BV650 (clone N418, Biolegend), Ly6C-PE (clone HK1.4, Biolegend), CD64-APC (clone X54-5/7.1, Biolegend), CD11c-BV570 (clone M1/70, Biolegend), and CX3CR1-BV510 (clone SA011F11, Biolegend) antibodies and the LD FVS700 fixable dye (cat# 564997, BD Biosciences). After staining, cells were washed with 1 mL of flow cytometry buffer, and fixed in 1x IC fixation buffer (e-biosciences) for 20 min at RT in the dark. Fixed cells were washed and resuspended in flow cytometry buffer, and acquired on a LSRII flow cytometer (BD Biosciences). Unstained cells and FMOs samples were also acquired as controls. Data was acquired with the FACSDiva software (BD Biosciences) and analyzed with FlowJo software (Treestar Inc).

### RNA isolation

Total RNA from brain tissue and BECs were isolated by TRIzol method according to the manufacturer’s instructions (Thermo Fisher Scientific). For brains tissue and cells, 1ml of TRIzol was used to homogenize the tissue by passing it through a syringe several times. The lysates were transferred to Phase Lock Gel 2ml tubes, and 200µl of chloroform (Thermo Fisher Scientific) was added to each tube, mixing vigorously for 15 seconds, followed by incubation at room temperature for 3 minutes. Samples were then centrifuged at 12000g for 10 minutes at 4°C, and the aqueous phases containing RNA were collected and transferred to 1.5ml DNAse/RNAse free microfuge tubes. To precipitate RNA, 500µl of isopropanol was added, samples were resuspended and incubated for 10 minutes at room temperature followed by centrifugation at 12000g for 10 minutes at 4°C. The supernatant was removed and the pellet was washed with 1ml of 75% ethanol followed by centrifugation at 7500g for 5 minutes at 4°C. RNA was resuspended in water and the quantity (ND-1000 spectrophotometer; NanoDrop Technologies) and quality (TapeStation; Agilent) of total RNA were analyzed.

### RT-qPCR analysis

A total RNA amount of 300 ng was used to produce single-stranded complementary DNA (cDNA) using random primers according to the manufacturer’s protocol (Thermo Fisher Scientific). For gene expression analysis, 10 ng of cDNA was used with the Kapa SybrFast qPCR master mix (Kapa Biosystems) and thermal cycler (CFX96 Real-Time System; Bio-Rad) were used to determine the relative levels of the genes analyzed according to the manufacturer’s protocol. Actin mRNA levels were used as internal control, and the 2^-ΔΔCT^ method was used for analyses of the data ^10^.

### RNA-sequencing (RNA-seq)

Total RNA was assessed for quality using an Agilent Tapestation 4200, and 50 nanograms of RNA from samples with an RNA Integrity Number (RIN) greater than 8.0 were used to generate RNA-seq libraries using the Illumina® Stranded mRNA Prep (Illumina, San Diego, CA). Samples were processed following manufacturer’s instructions. Resulting libraries were multiplexed and sequenced with 100 basepair (bp) Paired End reads (PE100) to a depth of approximately 25 million reads per sample on an Illumina NovaSeq 6000. Samples were demultiplexed using bcl2fastq Conversion Software (Illumina, San Diego, CA). Sequencing analysis was performed using the R programming environment and the RiboSeq systemPipeR workflow. All reads were aligned to the Mus Musculus GRCm39 genome version 104 using hisat2 read alignment software. Read counting was performed using the GenomicFeatures library and the corresponding GRCm39 GTF file from ensemble. All fold change calculations were performed using EdgeR.

### Inflammasome activity assay

BECs were isolated and resuspended in EBM-2 media (Lonza) supplemented with 0.1% gentamicin, 0.1% ascorbic acid, 0.1% heparin, and 0.2% BSA. Cells were incubated with the NLRP3 inhibitor MCC950 Sodium (10 µM;) or vehicle and seeded onto poly-lysine-treated coverslips for 1 hour at 37°C. After incubation, coverslips were washed three times using 1X HBSS + Ca^+2^, and then incubated with 30X FAM-FLICA Caspase-1 probe (ImmunoChemistry Technologies) in EBM-2 media pluss supplements for 1 hour at 37°C as described by the manufacturer’s protocol (ImmunoChemistry Technologies). Rat monoclonal antibody against VCAM1 (Alexa-674 labelled, clone 429(MVCAM.A)) and CD45 (Alexa-594, clone S18009D) were incubated at room temperature for 30 min (BioLegend). Preparations were fixed in 4% PFA in PBS and mounted on microscope slides using Fluoromount-G mounting medium (SouthernBiotech). The slides were viewed with a fluorescent microscope (Keyence), and the images were captured with BZX-700 software (Keyence). The quantification analysis was performed using ImageJ Ver.1.53f on high-resolution images.

### Immunohistochemistry

Brains from *Pdcd10^BECKO^* and littermate control *Pdcd10^fl/fl^* mice at the specific age points were isolated and fixed in 4% PFA in PBS at 4°C overnight. Tissue was placed into cryoprotective 30% sucrose solution in PBS, and then embedded and frozen in O.C.T compound (Fischer Scientific). Brains were cut using a cryostat into 18-µm sagittal sections onto Superfrost Plus slides (VWE International). Sections were incubated in a blocking-permeabilization solution (0.5% Triton X-100, 5% goat serum, 0.5% BSA, in PBS) for 2 hours and then incubated in rabbit polyclonal antibodies against GFAP (1:300; GA524; Agilent Dako), rat polyclonal antibodies against GFAP (1:200; Thermo Fisher Scientific), rabbit polyclonal antibodies against Iba1 (1:100; 019-19741; FUJIFILM Wako) in PBS at room temperature overnight. Slides were washed one time in brain Pblec buffer (1X PBS, 1mM CaCl2, 1mM MgCl2, 0.1 mM MnCl2, and 0.1% Triton X-100) and incubated with isolectin B4 (IB4) FITC conjugated (1:80, L2895; Sigma-Aldrich) in brain-Pblec buffer at 4°C overnight. Tissue sections were washed four times in PBS and incubated with suitable Alexa Fluor coupled secondary antibodies (1:300, Thermo Fisher Scientific) in PBS for 2h at RT. Cell nuclei were stained with DAPI and mounted using Fluoromount-G mounting medium (SouthernBiotech). Fresh frozen brain tissue was used to perform immunofluorescence analysis of leukocyte infiltration. Rat monoclonal antibody against CD41 (Alexa-488/594, clone MWReg30), CD45 (Alexa-fluor 594, clone S18009D), CX3CR1 (Alexa-fluor 488, clone SA011F11), LY6G (Alexa-fluor 488, clone 1A8), CD16/32 (PE/Dazzle 594, clone S17011E), CD206 (Alexa-fluor 488, clone C068C2), were incubated at room temperature for 30 min (BioLegend). Preparations were fixed in 4% PFA in PBS and incubated with isolectin B4 (IB4) biotin conjugated (1:80) in brain-Pblec buffer at 4°C overnight. After streptavidin (Alexa-Fluor 647) staining the cell nuclei were stained with DAPI and mounted with Fluoromount-G mounting medium (SouthernBiotech). The slides were viewed with a high-resolution slide scaner (Olympus VS200 Slide Scaner), and the images were captured with VS200 ASW V3.3 software (Olympus). Quantifications were performed blinded. The quantification analysis was performed using ImageJ Ver. 1.53f on high-resolution images. Four to five brain sections per mouse were used for analysis.

### CCM lesion quantification

Brains from *Pdcd10^BECKO^Ikkb^+/+^*, *Pdcd10^BECKO^*;*Ikkb^BECKO/+^* and *Pdcd10^BECKO^*;*Ikkb^BECKO^* mice were isolated and fixed in 4% PFA at 4°C overnight. After cryoprotection, sucrose (30%) and freezing, 18 µm sections of sagittal brain tissue were cut onto Superfrost Plus slides (VWR International), and stained by the hematoxylin and eosin (5 brain sections/mouse, sections were performed from superior sagittal sinus to the cerebral hemisphere). Slides were imaged using NanoZoomer Slide Scanner (Hamamatsu Photonics; San Diego, USA). Lesions were analyzed as stage 1, single cavernous, or stage 2, multi cavernous and thrombosis. Quantifications were performed blinded. The quantification analysis was performed using Hamamatsu Photonics software.

### Statistical analysis

Data are expressed as average values ± standard error of the mean (SEM) for multiple individual experiments. For all experiments, the number of independent and biological replicates (n) is indicated. The sample sizes were estimated with a 2-sample t test (2 tailed). A 2-tailed unpaired Student t test was used to determine statistical significance. For multiple comparisons, 1-way analysis of variance (ANOVA), followed by the Tukey post hoc test, was used.

## Acknowledgments

The authors thank Mark H. Ginsberg for helpful discussion; Yin Shi (Sherly), Cassandra Bui, Victoria Herrera and Wilma McLaughlin for technical assistance; Jennifer Santini for microscopy technical assistance; and Kristen Jepsen for RNA-seq technical assistance; *Ikkbfl/fl* mice were the generous gift of Michael Karin (UCSD);*Slco1c1-CreERT2 mice* were the generous gift of Markus Schwaninger (University of Lübeck); *Pdcd10^fl/fl^* mice were the generous gift of Wang Min (Yale University).

